# TRAMP assembly alters the conformation and RNA binding of Mtr4 and Trf4-Air2

**DOI:** 10.1101/2024.07.25.605035

**Authors:** Joshua M. Denson, Naifu Zhang, Darby Ball, Kayla Thompson, Sean J. Johnson, Sheena D’Arcy

## Abstract

The TRAMP complex contains two enzymatic activities essential for RNA processing upstream of the nuclear exosome. Within TRAMP, RNA is 3’ polyadenylated by a sub-complex of Trf4/5 and Air1/2 and unwound 3’ to 5’ by Mtr4, a DExH helicase. The molecular mechanisms of TRAMP assembly and RNA shuffling between the two TRAMP catalytic sites are poorly understood. Here, we report solution hydrogen-deuterium exchange data with thermodynamic and functional assays to uncover these mechanisms for yeast TRAMP with Trf4 and Air2 homologs. We show that TRAMP assembly constrains RNA-recognition motifs that are peripheral to catalytic sites. These include the Mtr4 Arch and Air2 zinc knuckles 1, 2, and 3. While the Air2 Arch-interacting motif likely constrains the Mtr4 Arch via transient interactions, these do not fully account for the importance of the Mtr4 Arch in TRAMP assembly. We further show that tRNA binding by single active-site subunits, Mtr4 and Trf4-Air2, differs from the double active-site TRAMP. TRAMP has reduced tRNA binding on the Mtr4 Fist and RecA2 domains, offset by increased tRNA binding on Air2 zinc knuckles 2 and 3. Competition between these RNA-binding sites may drive tRNA transfer between TRAMP subunits. We identify dynamic changes upon TRAMP assembly and RNA-recognition motifs that transfer RNA between TRAMP catalytic sites.

## Introduction

RNA surveillance pathways in eukaryotes ensure the proper quality and abundance of coding and non-coding RNAs (1-3). Impaired RNA surveillance mechanisms cause pathologies such as immune disorders, neurodegenerative diseases, and various cancers (4-7). The exosome is a macromolecular protein complex with endo- and exo-nuclease activity central to RNA processing and decay (8-11). The TRAMP complex is a conserved exosome co-factor that modifies and delivers targeted RNA substrates for decay (12-17). In *Saccharomyces cerevisiae*, the TRAMP complex contains poly(A) polymerase Trf4 or 5, RNA-binding protein Air1 or 2, and ATP-dependent RNA helicase Mtr4. The two enzyme activities of the TRAMP complex ensure that an RNA substrate is polyadenylated at its 3’ end and actively unwound before entering the exosome (18). The combination of Trf4/5 and Air1/2 homologs in TRAMP influences RNA substrate selectivity. Here, we focus on TRAMP containing Trf4 and Air2, which preferentially targets RNAP III transcripts such as aberrant tRNAs (19). We investigate interfaces and conformational changes associated with TRAMP assembly and compare tRNA-binding of single active-site subunits, Mtr4 and Trf4-Air2, and the fully-assembled TRAMP complex.

Mtr4 is an essential 3’ to 5’ ATP-dependent RNA helicase that contains five domains (**Fig. S1A**) (20). Four domains, the RecA1, RecA2, Winged-Helix, and Helical Bundle, comprise the helicase core that contacts RNA through well-characterized motifs (9, 21, 22). The fifth domain, the Arch, is an insertion in the Winged-Helix and contains a helical stalk and globular Fist or KOW region (23-25). Arch positioning relative to the helicase core is dynamic and modulated by interactions with RNA or regulatory proteins (21, 26). Several proteins bind to the Fist through a short linear sequence called an Arch-Interacting Motif (AIM) (21, 27-31). In the context of TRAMP, Air2 may use an AIM spanning residues 7-14 in its disordered N-terminal region to interact with the Mtr4 Fist. This hypothesis and our current molecular understanding of TRAMP assembly come from a crystal structure of Mtr4 with an engineered chimera of Air2 residues 1 to 62 (Air2_1-62_) and Trf4 residues 111 to 160 (Trf4_111-160_) (**Fig. S1A**). Within the crystal lattice, a single Air2_5-52_ interacts with two Mtr4, one via the AIM binding to the Fist and the other via Air_20-52_ binding on the surface of the helicase core (31). The distance between the helicase core and the Fist of a single Mtr4 is too large to accommodate Air2_5-52_ binding at both sites unless there is significant repositioning of the Arch. It remains uncertain whether a single copy of Mtr4 and Air2 can simultaneously engage at both interaction sites and how such an interaction would influence Arch dynamics.

Downstream of the AIM and helicase core binding sites, Air2 contains five loosely structured CX_2_CX_4_HX_4_C zinc knuckles (ZnKs 1-5) (**Fig. S3A**) (32). These ZnKs mediate binding to Trf4, as well as RNA. Trf4 belongs to the Pol β family of nucleotidyl-transferases and adds adenosine nucleotides to free, unpaired, 3’ RNA ends with at least one nucleotide overhang (33-35). The Trf4 catalytic domain is inserted within a central domain that contains a type 2 Nucleotide Recognition Motif (NRM) (36). The interaction between Trf4 and Air2 is very tight and critical for polymerase activity of recombinant Trf4 (16, 17, 37). It involves Air2 ZnKs 4-5 sitting on the Trf4 central domain (**Fig. S4B**) (36). The precise conformation and positioning of Air2 ZnKs 1-3 in the Trf4-Air2 complex is unclear and is of interest as ZnKs 2-4 may directly bind RNA. NMR studies show that adding hypomodified tRNA_i_^Met^ to Air2 without Trf4 alters ZnKs 2-4 (32). There is a clear need to understand the molecular details of RNA recognition by Trf4-Air2 as a sub-complex and within TRAMP.

TRAMP also contains an interface between Mtr4 and Trf4, albeit a small one, as Trf4_120-127_ forms a short β-strand that extends the Mtr4 RecA2 β-sheet (**Fig. S1A**) (31). This small segment of Trf4 is within the ∼150-residue N-terminal region of Trf4 that is flexible in solution (36). The disordered nature of the Mtr4 recognition sites of both Trf4 and Air2 makes it challenging to predict the relative orientation and proximity of the two TRAMP active sites and, thus, their crosstalk mechanism. TRAMP assembly enhances Mtr4 helicase ∼four-fold and causes Trf4 to pause after adding 5-7 adenosine nucleotides, making the RNA an ideal substrate for Mtr4 (35, 38). Presumably, RNA substrates are first modified by Trf4, then transferred to Mtr4 to be unwound and fed into the exosome. It is mechanistically unclear how the subunits of TRAMP impact each other’s activities and how RNA substrates transfer between the two active sites.

We have used hydrogen-deuterium exchange with mass spectrometry (HDX) and complementary biochemical assays to address gaps in understanding TRAMP assembly and RNA binding. We demonstrate that TRAMP assembly imposes constraints on the Mtr4 Arch and Air2 ZnKs 1-3, which are involved in tRNA recognition. We map the tRNA binding site on the Trf4-Air2 sub- complex and provide a compelling model that includes all Air2 ZnKs. These data provide the foundation for directly comparing tRNA binding to double active-site TRAMP and single active-site Mtr4 or Trf4-Air2 sub-complexes. We uncover apparent competition for tRNA between the Mtr4 Fist and RecA2 domains and Air2 ZnKs 2-3, suggesting they are involved in tRNA transfer within TRAMP. Our solution analysis enhances the mechanistic understanding of TRAMP assembly and tRNA recognition and transfer.

## Results

### Solution interactions between the Mtr4 helicase core and Trf4-Air2

To investigate interfaces involved in TRAMP assembly in solution, we performed HDX. We compared TRAMP’s deuterium uptake (D-uptake) to Mtr4 alone or a co-expressed Trf4-Air2 complex. We used full-length Mtr4, a catalytically dead point mutant of full-length Trf4 (D236, 238A), and a C-terminally truncated Air2 (Air2_1-223_) that includes the proposed AIM (residues 7- 14) and all five ZnKs. Differences in D-uptake indicate altered hydrogen bond stability of amides in the protein backbone due to either protein interaction or conformational change (39). We monitored peptides covering 90% of Mtr4, 84% of Trf4, and 100% of Air2_1-223_ (**Table S1**). For Mtr4, the HDX data were consistent with the prior TRAMP structure (**Fig. S1A-B**), especially in the RecA2 (**Fig. 1A, E**) and Helical Bundle domains (**Fig. S1C, E**) of the helicase core. We saw a reduction in D-uptake for the outermost strand of the Mtr4 RecA2 β-sheet and nearby surfaces, as well as for the surface formed by the Mtr4 RecA2 and Helical Bundle domains (**Fig. 1C**). The reduction in D-uptake was specifically located at Trf4-Air2 binding surfaces as non-binding RecA1 and Winged-Helix domains showed no change in D-uptake upon TRAMP formation (**Fig. 1A, D**). Correspondingly, eight of nine peptides covering Air2_21-52_ (**Fig. 3A, D**) and one of five peptides covering Trf4_120-127_ (**Fig. 4A**) showed a significant reduction in D-uptake upon TRAMP formation. We do not have coverage of Trf4 isoleucine 124, which forms a primary contact in the hybrid Trf4- Mtr4 β-sheet. Beyond Trf4_120-127_, we saw a slight reduction in D-uptake around the Trf4 catalytic pocket and a slight increase in D-uptake around the nucleotide recognition motif (**Fig. 4A**). The minimal impact of TRAMP formation on Trf4_120-127_, as well as on the entire Trf4 protein, reflects the small size of the interface between Trf4_120-127_ and Mtr4, and shows that any additional direct interactions must be weak or transient in solution. Although we used longer Trf4 and Air2 constructs in our HDX, the interfaces we identified between Trf4-Air2_1-223_ and the Mtr4 helicase core match prior crystallographic analysis.

**Figure 1.**
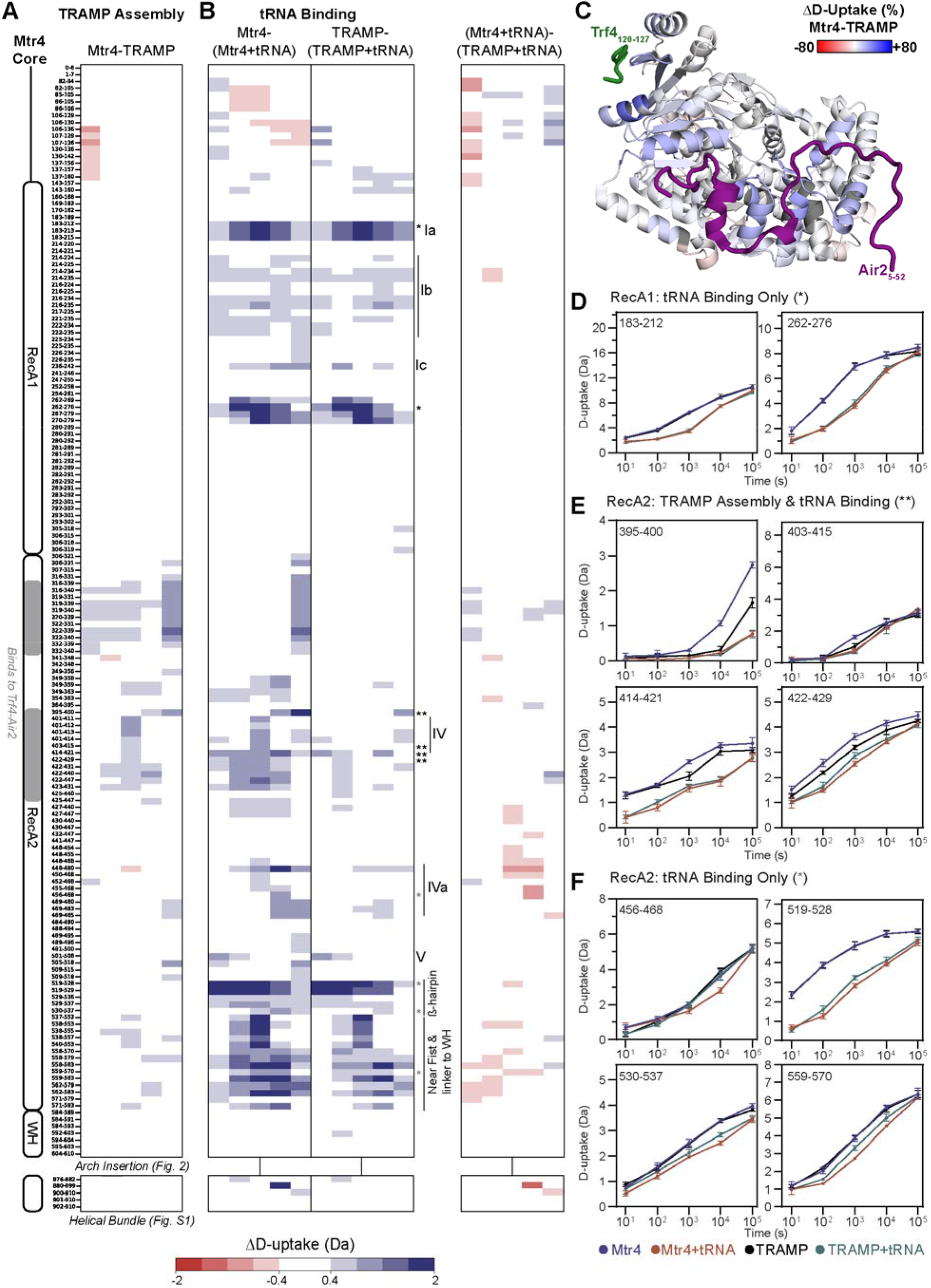
The Mtr4 helicase core is involved in TRAMP assembly and tRNA binding. (**A**) The difference in Mtr4 helicase core D-uptake upon TRAMP assembly. (**B**) The difference in Mtr4 helicase core D-uptake upon tRNA binding to Mtr4 (left) or TRAMP (middle); direct comparison of the two tRNA-bound complexes (right). For **A** and **B**, red/blue indicates a difference of ≥|0.4| Da and a *p*-value ≤0.01 in a one-sided Welch’s t-test (n=3). Exchange time points were 10^1^-10^5^ s (y-axis), and the Mtr4 schematic is based on PDB 4U4C. (**C**) Mtr4 helicase core colored by the difference in percent D-uptake per residue between Mtr4 and TRAMP summed across all time points. The scale is -80 to 80% (red to white to blue) based on DynamX residue-level scripts without statistical filtering. Mtr4 residues without coverage are gray, Trf4 is green, and Air2 is purple. The full TRAMP structure is shown in **Fig. S1A-B**. The structure is PDB 4U4C. (**D**) Select D-uptake plots of Mtr4 RecA1 peptides involved in tRNA binding (*). (**E, F**) Select D-uptake plots of Mtr4 RecA2 involved in TRAMP assembly and tRNA binding (**) or only tRNA binding (gray*). For **D-F**, D-uptake plots show Mtr4 (blue), Mtr4 with tRNA (orange), TRAMP (black), and TRAMP with tRNA (green). Error bars are ±2SD of the average with n=3. The y-axis is 80% of the maximum theoretical D-uptake, assuming complete back exchange of the N-terminal residue of each peptide. Similar data for the Mtr4 Helical Bundle are in **Fig. S1C-E**.

**Figure 2.**
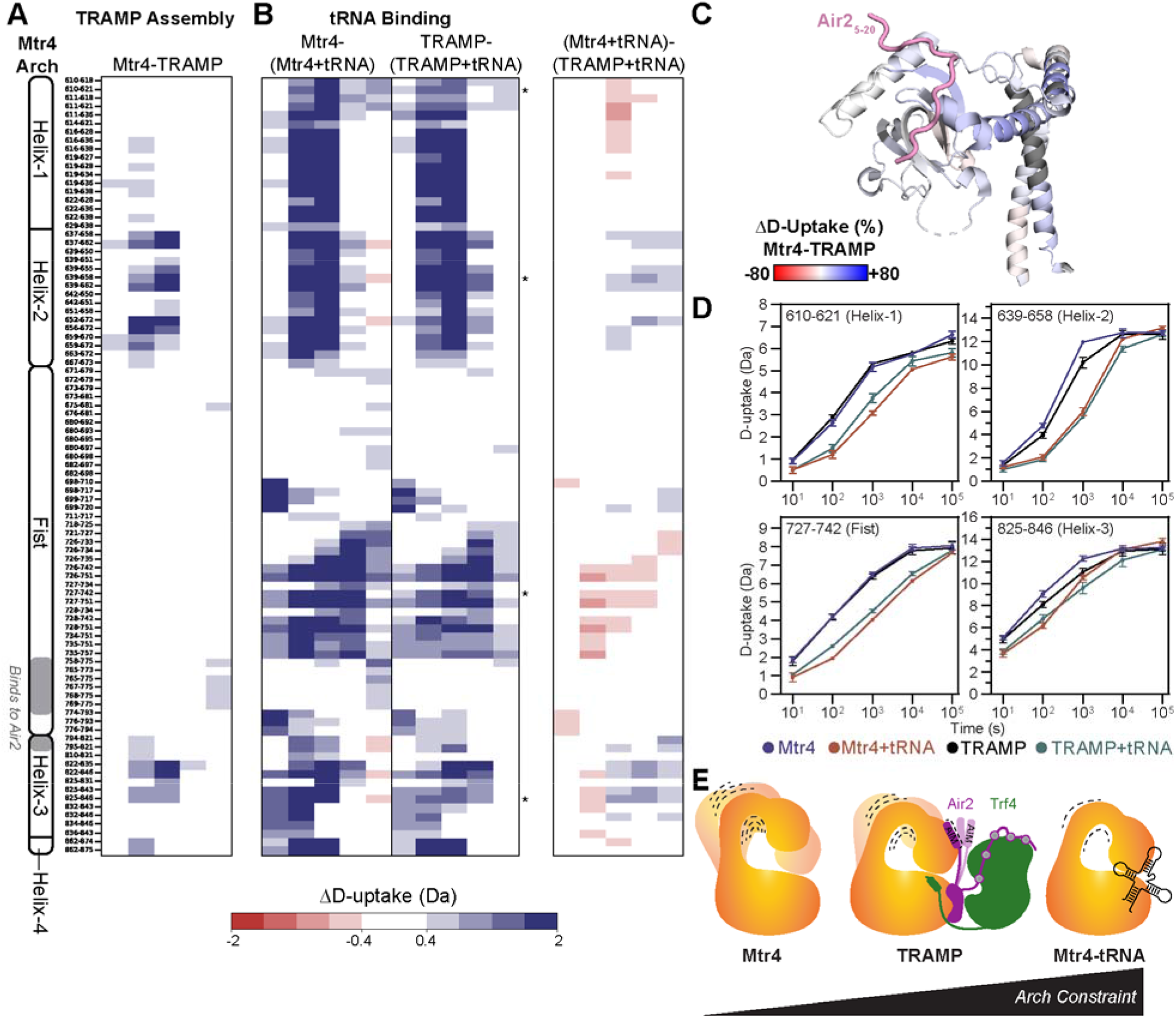
TRAMP assembly and tRNA binding alter the constraints on the Mtr4 Arch. (**A**) The difference in Mtr4 Arch D-uptake upon TRAMP assembly. (**B**) The difference in Mtr4 Arch D-uptake upon tRNA binding to Mtr4 (left) or TRAMP (middle); direct comparison of the two tRNA-bound complexes (right). For **A** and **B**, red/blue indicates a difference of ≥|0.4| Da and a *p*- value ≤0.01 in a one-sided Welch’s t-test (n=3). Exchange time points were 10^1^-10^5^ s (y-axis), and the Mtr4 schematic is based on PDB 4U4C. (**C**) Mtr4 Arch colored by the percent difference in D-uptake per residue between Mtr4 and TRAMP summed across all time points. The scale is - 80 to 80% (red to white to blue) based on DynamX residue-level scripts without statistical filtering. Mtr4 residues without coverage are gray, and Air2 from a symmetry-related Mtr4 is pink. The full TRAMP structure is shown in **Fig. S1A-B**. The structure is PDB 4U4C. (**D**) Select D-uptake plots of Mtr4 Arch peptides with Mtr4 (blue), Mtr4 with tRNA (orange), TRAMP (black), and TRAMP with tRNA (green). Error bars are ±2SD of the average with n=3. The y-axis is 80% of the maximum theoretical D-uptake, assuming complete back exchange of the N-terminal residue of each peptide. (**E**) Relative constraint of the Mtr4 Arch in various complexes. The Arch becomes constrained in TRAMP, likely due to transient interactions between the Mtr4 Fist and Air2 AIM. The Arch is most constrained upon tRNA binding.

**Figure 3.**
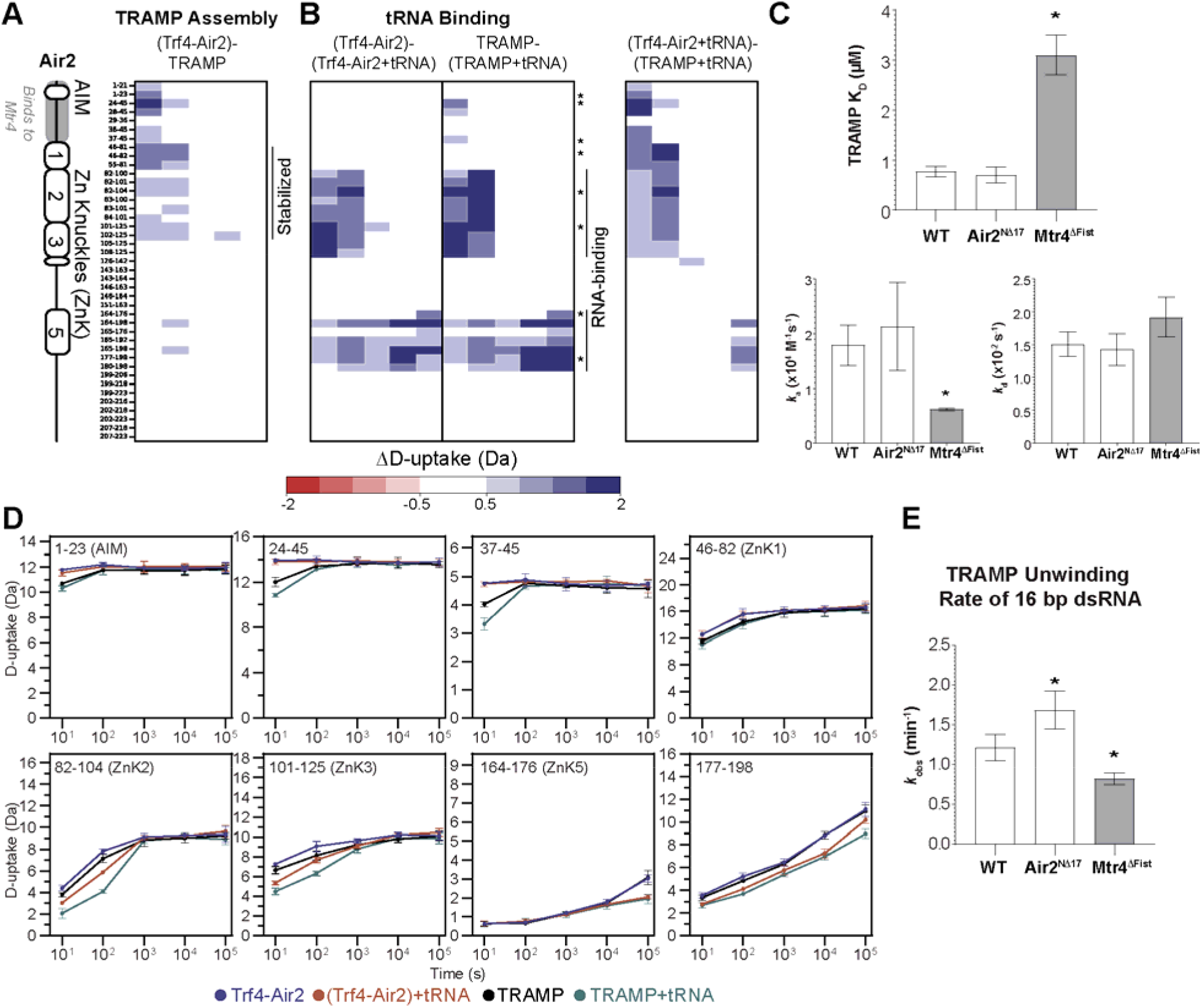
Air2 has a multi-faceted role in TRAMP assembly and tRNA binding. (**A**)The difference in Air2 D-uptake upon TRAMP assembly. (**B**) The difference in Air2 D-uptake upon tRNA binding to Trf4-Air2 (left) or TRAMP (middle); direct comparison of the two tRNA- bound complexes (right). For **A** and **B**, red/blue indicates a difference of ≥|0.5| Da and a *p*-value ≤0.01 in a one-sided Welch’s t-test (n=3). Exchange time points were 10^1^-10^5^ s (y-axis), and the Air2 schematic is based on PDB 2LLI. (**C**) TRAMP assembly parameters (K_D_, *k*_*a*_, and *k*_*d*_) measured by SPR with Mtr4 in solution and Trf4-Air2 immobilized via nickel binding of an N- terminal hexa-histidine tag on Air2. Representative response curves and complementary SEC data are in **Fig. S2**. (**D**) Select D-uptake plots of Air2 peptides with Trf4-Air2 (blue), Trf4-Air2 with tRNA (orange), TRAMP (black), and TRAMP with tRNA (green). Error bars are ±2SD of the average with n=3. The y-axis is 80% of the maximum theoretical D-uptake, assuming complete back exchange of the N-terminal residue of each peptide. (**E**) Helicase activity using a synthetic 16 bp dsRNA substrate with a 5x adenosine 3’ overhang and 500 nM TRAMP. Representative gels and similar experiments with a 32 bp dsRNA are in **Fig. S3F-G**. For **C** and **E**, TRAMP was wild-type (WT), with Air2^NΔ17^, or with Mtr4^ΔFist^ (gray). An asterisk (*) indicates a *p*-value ≤0.05 in a one-way ANOVA with the posthoc Tukey HSD test. Error bars show ±SD of the average with n=3, including at least 1 biological replicate. Complementary data for Air2^ZKO^ are in **Fig. S3**.

**Figure 4.**
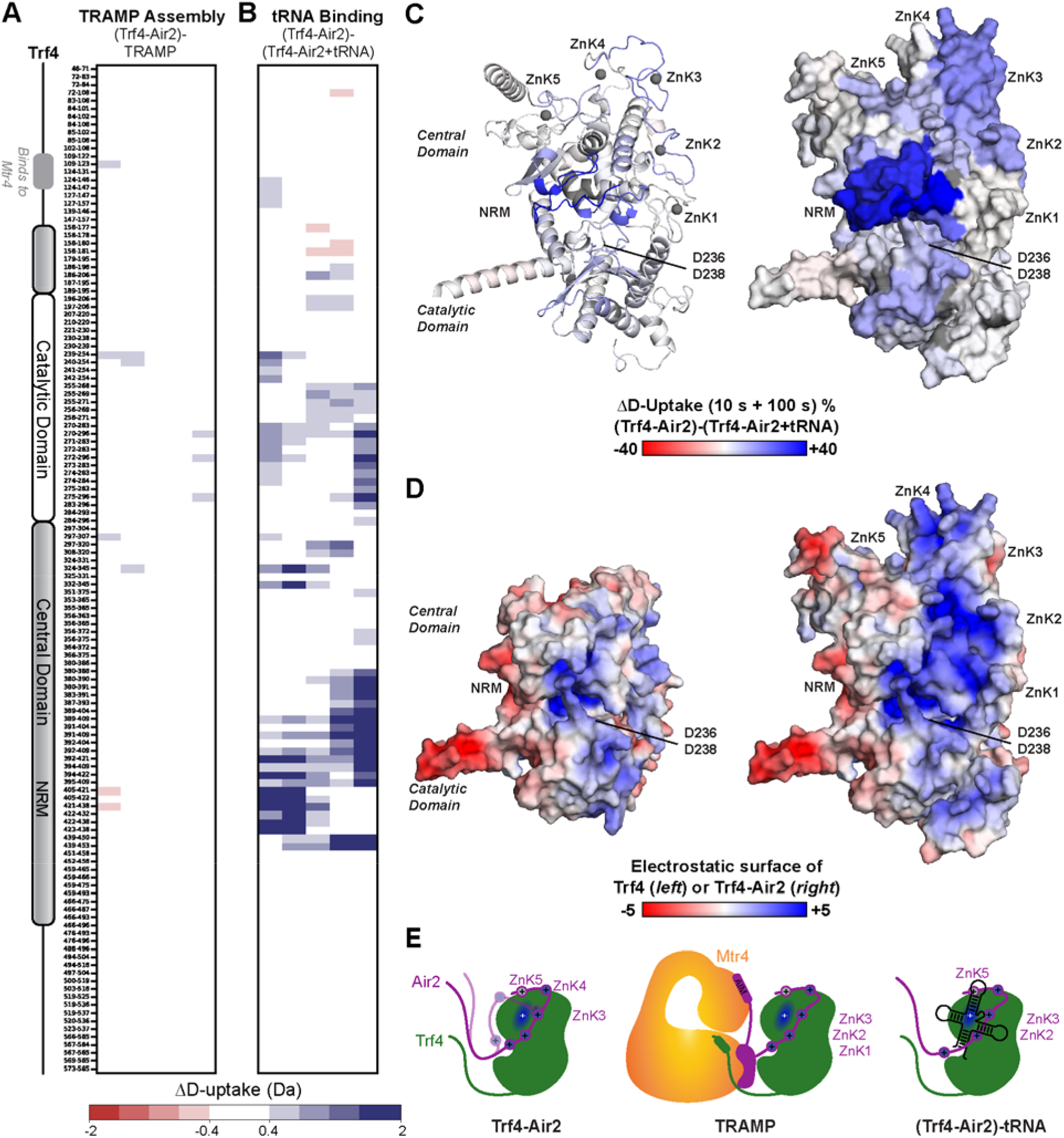
tRNA binds to a negative patch formed by Trf4 and Air2. (**A**) The difference in Trf4 D-uptake upon TRAMP assembly. (**B**) The difference in Trf4 D-uptake upon tRNA binding to Trf4-Air2. For **A** and **B**, red/blue indicates a difference of ≥|0.4| Da and a *p*- value ≤0.01 in a one-sided Welch’s t-test (n=3). Exchange time points were 10^1^-10^5^ s (y-axis), and the Trf4 schematic is based on PDB 3NYB. (**C**) Alphafold2 Multimer model of Trf4-Air2 colored by the percent change in D-uptake per residue upon adding tRNA summed across 10^1^ and 10^2^ s time points (left as cartoon, right as surface). The scale is -80 to 80% (red to white to blue) based on DynamX residue-level scripts without statistical filtering. Trf4-Air2 residues without coverage are gray, and zinc atoms are spheres. Zincs were placed in the model by pairwise superposition of coordinating residues in prior structures. (**D**) Alphafold2 Multimer model of Trf4 (left) and Trf4-Air2 (right) colored by electrostatic potential. The scale is -5 to 5 (red to white to blue) calculated by APBS (53). For **C** and **D**, images contain only high-confidence regions (pLDDT > 40) which are Trf4_133-523_ and Air2_24-220_ (see **Fig. S4A**). (**E**) Cartoon depicting the role of Air2 ZnKs in various complexes. Air2 ZnK3, ZnK4, and ZnK5 interact with Trf4. Air2 ZnK1, ZnK2, and ZnK3 become ordered upon TRAMP assembly and form a continuous negative patch with Trf4. Air2 ZnK2, ZnK3, and ZnK5 are altered upon tRNA binding either from direct interactions or conformational change. Additional data looking at tRNA binding by Trf4-Air2 or TRAMP are in **Fig. S4**.

### TRAMP assembly constrains the dynamics of the Mtr4 Arch

Beyond the Mtr4 helicase core, we also observed that TRAMP formation caused a reduction in D-uptake in the Mtr4 Arch (**Fig. 2A**). The Arch is composed of a stalk with four helices (helices 1-4) and a Fist domain between helices 2 and 3 (**Fig. S1A-B**). TRAMP formation reduced D-uptake in helices 2 and 3, with a more subtle reduction in helices 1 and 4, and essentially no change in D-uptake in the Fist (**Fig. 2A, C, D**). The reduced D-uptake suggests that TRAMP formation constrains the conformational dynamics of the Mtr4 Arch. We reported a similar result for Mtr4 binding to tRNA and showed that direct contact between the Fist and tRNA drives the Arch constraint (26). We repeated these latter findings by examining Mtr4 and Mtr4 with tRNA (1:2 molar ratio) parallel to TRAMP samples (**Fig. 2B left panel**). We obtained the same result as previously, with an additional discovery that Mtr4 residues 699-717, which comprise a loop in the Fist, also bind tRNA. Peptides covering these residues were not found in our prior experiment. Direct comparison of the effect of TRAMP formation to the effect of tRNA binding clearly shows a greater reduction in Arch dynamics for tRNA binding (compare **Fig. 2A** and **Fig. 2B left panel**). The D-uptake difference in the Arch helical stalk is greater and more widespread for tRNA binding than for TRAMP formation. The relatively similar pattern of reduced D-uptake over time across stalk peptides, however, does suggest that the conformation of the constrained Arch is similar between TRAMP and tRNA-bound Mtr4. Compared to Mtr4 alone, the constrained conformation of the Mtr4 Arch occurs more frequently upon TRAMP formation, albeit not as frequently as when the Fist directly binds tRNA (**Fig. 2E**).

A possible mechanism for Mtr4 Arch constraint in TRAMP is direct contact between the Air2 AIM (residues 7-14) and the Mtr4 Fist. This interface occurs in the TRAMP structure via crystal contacts; residues 5 to 20 of Air2_1-62_ bind to the Fist of one Mtr4, while residues 21-52 of Air2_1-62_ bind to the helicase core of another Mtr4 (**Fig. S1A**). Given that TRAMP does not oligomerize (31), it is feasible that a single Air2 could simultaneously bind the Mtr4 Fist and helicase core. The HDX data suggest that the interface between the Air2 AIM and Mtr4 Fist does occur but is very weak and transient in solution. We observed reduced D-uptake on the Fist surface upon TRAMP formation, but the reduction was too small to pass our stringent heatmap thresholds (**Fig. 2A**). The reduction is evident only when the summed difference across all time points is projected onto Mtr4 with no statistical filtering (**Fig. 2C**). The location of the reduction maps nicely to the binding site of the Air2 AIM seen in the TRAMP structure. From the Air2 perspective, we also saw reduced D-uptake upon TRAMP formation for the two AIM-containing peptides (**Fig. 3A, D**). These peptides, however, are long (residues 1-21 and 1-23) and slightly overlap with the start of the interface between Air2 and the Mtr4 helicase core.

We used two deletion mutants to investigate further the interaction between the Air2 AIM and the Mtr4 Fist. The first was Mtr4 with the Fist domain replaced by three glycine residues (Mtr4^ΔFist^), and the second was Air2_1-223_ lacking residues 1-17 (Air2^NΔ17^). Air2^NΔ17^ removes the AIM but retains all residues critical for Air2 interaction with the Mtr4 helicase core. We assayed the ability of these mutants to form TRAMP using size exclusion chromatography and surface plasmon resonance (SPR) with Trf4-Air2 immobilized via an N-terminal his-tag (**Fig. S2**). We saw that Trf4-Air2^NΔ17^ bound Mtr4 with similar efficiency to wild-type and that the resulting complex had the same oligomeric state (**Fig. S2A-B**). Thermodynamic and kinetic constants of Mtr4 binding were also equivalent for Trf4-Air2^NΔ17^ and wild-type (**Fig. 3C** and **Fig. S2C**). Finally, activity assays using TRAMP with 16-bp or 32-bp dsRNA substrates show that AIM deletion had little to no effect on unwinding (**Fig. 3E** and **Fig. S3F-G**). These data show that deletion of the Air2 AIM has no measurable impact on TRAMP assembly or unwinding activity, consistent with only weak or transient interactions with Mtr4 in solution.

In contrast, similar experiments with Mtr4^ΔFist^ showed inefficient TRAMP assembly, with the bulk of Mtr4^ΔFist^ failing to bind Trf4-Air2. This result was unexpected as previous work has demonstrated that Mtr4 can form a complex with Trf4-Air2 with the entire Arch domain removed (24, 40). We saw a clear population of unbound Mtr4^ΔFist^ when mixed with an equimolar amount of Trf4-Air2 (**Fig. S2A-B**). We also measured the binding affinity of Mtr4 to Trf4-Air_1-223_ to be four-fold weaker for Mtr4^ΔFist^ than for wild-type (∼3 μM versus ∼0.8 μM) (**Fig. 3C**). This reduction in affinity is due to a compromised association rate (6173.3 M^-1^s^-1^ versus 17866.7 M^-1^s^-1^) with an unchanged dissociation rate (**Fig. 3C** and **Fig. S2C**). Consistently, we found that deletion of the Fist compromised TRAMP unwinding of dsRNA substrates irrespective of the involvement of the Fist in RNA binding (**Fig. 3E** and **Fig. S3F-G**). The 16-bp dsRNA does not interact with the Fist, while the 32-bp dsRNA does (26). Altogether, these data identify a role for the Mtr4 Fist in TRAMP assembly beyond its binding to the Air2 AIM. TRAMP formation may require the Mtr4 Arch to visit a less dynamic conformation likely driven by intramolecular interactions between the Fist and helicase core. A weak or transient interface between the Air2 AIM and Mtr4 Fist could play a secondary role once TRAMP has assembled.

### TRAMP assembly constrains the conformation of Air2 ZnKs 1-3

A final observation in the HDX data is that TRAMP assembly reduced the D-uptake of Air2 ZnKs 1-3. These ZnKs were not included in the partial structure of TRAMP (31) (**Fig. S1A**) or Trf4-Air2 (36) (**Fig. S4B**) but were visualized by NMR of Air2 alone (32) (**Fig. S3A**). Reduced D-uptake occurred upon TRAMP formation in several overlapping peptides at both the first and the second time points of exchange (**Fig. 3A, D**). This reduction could be due to new Air2 interfaces with Mtr4 or Trf4 in TRAMP (for which there is no precedent), or a conformational change in Air2. To assess these possibilities, we made a double point mutant of Air2 (C84S, C104S) that abolishes zinc coordination in ZnKs 2-3 (Air2^ZKO^). We characterized the effect of these mutations by comparing the D-uptake of wild-type Trf4-Air2 to mutant Trf4-Air2^ZKO^ (**Fig. S3B**). The mutations caused a dramatic increase in D-uptake for Air2 peptides covering ZnKs 1-3 (**Fig. S3B left panel**), with no change observed for the rest of Air2 or Trf4 (**Table S1**). The Air2 ZnK 1-3 peptides from the mutant reached maximal D-uptake by the 10 s time point, indicating a loss of structure for ZnKs 2 and 3 as expected, but also for ZnK 1. The apparent dependence of ZnK 1 integrity on the subsequent ZnKs shows cooperativity in Air2 folding within the Trf4-Air2 complex. The introduced mutations thus destabilized Air2 ZnKs 1-3 and did not alter Air2 interaction with Trf4.

We next compared TRAMP assembly and unwinding activity between wild-type Trf4-Air2 and mutant Trf4-Air2^ZKO^complexes. Using HDX, we saw a minimal effect, with almost all the changes in D-uptake described for wild-type TRAMP assembly also observed for the mutant (**Fig. S3C**). Similar changes occurred for Trf4 and Mtr4 peptides (**Table S1**) and Air2 outside of ZnKs 1-3 (**Fig. S3C**). For ZnKs 1-3, TRAMP formation did not reduce D-uptake in the mutant, as the three ZnKs remained destabilized as in Trf4-Air2^ZKO^ alone. TRAMP formation did not overcome the local destabilization of the three ZnKs caused by the mutations. SPR consistently measured similar thermodynamic and kinetic constants for Mtr4 binding to Trf4-Air2 or mutant Trf4-Air2^ZKO^ (**Fig. S3C-D**). Unwinding assays also showed identical activity toward a 16-bp dsRNA substrate for TRAMP with Trf4-Air2 or mutant Trf4-Air2^ZKO^ (**Fig. S3E-F**). These data ultimately show that the interactions between Mtr4, Trf4, and Air2 are unaffected by the loss of structure of Air2 ZnKs 1-3. We can thus conclude that the observed reduction in D-uptake of Air2 ZnKs 1-3 upon wild-type TRAMP assembly is due to a conformational change rather than a direct binding event (**Fig. 4E**). Conformational change of Air2 ZnKs 2-3 is of note as they mediate binding to RNA in Air2 alone (32).

### Trf4 and Air2 combine to form a positively charged tRNA-binding surface

Our HDX experiment also included Mtr4, Trf4-Air2_1-223_, and TRAMP, each with a 2-fold molar excess of hypomodified tRNA _i_ ^Met^ (**Table S1**). We can identify initial binding events, as the samples do not contain ATP, by comparing the D-uptake of proteins with and without tRNA. We have done this for Mtr4 [see above and (26)] and now do this for Trf4-Air2_1-223_. To best interpret the data, we have used the high-confidence regions (pLDDT>40) of an AlphaFold2 Multimer model of full-length Trf4-Air2 (41, 42) (**Fig. S4A**). The high-confidence regions encompass most residues in Trf4_133-523_ and Air2_24-220_ and superpose well with the structure of the Trf4 core and Air2 ZnKs 4-5 (RMSD 0.9 Å) (**Fig. S4B**). The model predicts two novel Trf4 helices and positions Air2 ZnKs 1-3 along the central domain of Trf4. The novel Trf4 helices are likely transient in solution as their peptides reached maximal D-uptake after only 10 s exchange. The D-uptake of the surface beneath Air2 after 10^4^ s exchange suggests stable contacts between Trf4 with ZnKs 3-5 but weak contacts with ZnKs 1-2 (**Fig. S4C**). The Trf4-Air2 model is valuable for visualizing the position of all Air2 ZnKs in the complex.

The presence of tRNA reduced D-uptake for specific regions of Trf4 (**Fig. 4B**) and Air2 (**Fig. 3B left panel**), showing that both proteins contribute to tRNA recognition. We observed distinct kinetic signatures with peptides having a change in either early time points (10 or 10^2^ s), late time points (10^4^ or 10^5^ s), or, in some cases, a combination of both with little or no change seen at the middle time point (10^3^ s). For Air2, ZnKs 2-3 showed reduced D-uptake early, while ZnK 5 showed reduced D-uptake late (**Fig 3B, D**). Similarly, for the Trf4 central domain, the NRM showed reduced D-uptake early, while the region upstream of the NRM showed reduced D-uptake late (**Fig. 4B**). The Trf4 catalytic domain showed a combination signature. These distinct kinetic signatures imply differing roles in tRNA recognition. Early changes are associated with direct binding, while late changes are related to conformational events or enhanced stability of the bound complex. For Trf4-Air2, the late changes localize to the known interface between the Trf4 central domain and Air2 ZnKs 4-5, consistent with tRNA enhancing the stability of this interface (**Fig. S4D**).

Mapping early reductions in D-uptake onto the model identifies the tRNA-binding site formed by Trf4 and Air2 (**Fig. 4C**). Changes map to the cleft between the Trf4 central and catalytic domains that house the NRM and catalytic residues (D236 and D238) respectively. This distribution of reduced D-uptake suggests Trf4 RNA recognition is similar to that in homologous proteins, as seen in the structure of homolog Tut7 with dsRNA (**Fig. S4E**) (43). The larger reduction for the NRM compared to the catalytic domain suggests it is the primary tRNA-recognition motif and that the active site does not stably bind our tRNA construct. Changes also map to Air2 ZnKs 2-3 that are proximal to the Trf4 NRM and bind tRNA in Air2 alone (**Fig. 4C**) (32). This distribution of reduced D-uptake confirms that these ZnKs maintain tRNA binding in Trf4-Air2. Air2 ZnKs 2-3 and the Trf4 NRM and catalytic domains must thus work together to bind and orient tRNA. The idea of a tRNA-binding surface formed by both Trf4 and Air2 is further suggested by the electrostatic potential of the Trf4-Air2 model (**Fig. 4D**). The model contains a continuous, positively charged surface that spans the Trf4 catalytic domain and NRM, and Air2 ZnKs 1-3. This surface is prime for interaction with the RNA backbone and closely mimics the distribution of reduced D-uptake upon adding tRNA. The Air2 ZnKs thus play a role in the Trf4-Air2 interface (ZnKs 3-5), TRAMP assembly (ZnKs 1-3), and Trf4-Air2 interaction with tRNA (ZnKs 2-3) (**Fig. 4E**).

### The binding of tRNA to Mtr4 or Trf4-Air2 is altered in TRAMP

Next, we compared the effect of tRNA on Mtr4 or Trf4-Air2 to the effect of tRNA on TRAMP. If tRNA binds to the subunits and TRAMP at the same location with similar thermodynamics, the difference map comparing the tRNA-bound subunits to tRNA-bound TRAMP will recapitulate the difference map for TRAMP assembly (**Fig. 1B, S1D, 2B, 3B**, and **S4F**). We observed that tRNA binding was virtually unchanged in TRAMP for Trf4 and the Mtr4 RecA1. We saw almost no difference in D-uptake between tRNA-bound Trf4 in Trf4-Air2 and TRAMP (**Fig. S4F**), as well as between tRNA-bound RecA1 in Mtr4 and TRAMP (**Fig. 1B**). This may seem surprising as Trf4 and the Mtr4 RecA1 both contain catalytic residues that act on the 3’ overhang of a tRNA substrate. Several explanations may account for this result. The tRNA may bind Trf4 and RecA1 simultaneously, which is feasible as RecA1 binds the end of the 3’ overhang that does not stably bind Trf4 (35), and the 3’ overhang in our tRNA construct is also quite long at 12 bases. Alternatively, in TRAMP, the tRNA may be transferred between Trf4 and the Mtr4 RecA1 rather than added to the free tRNA pool, as occurs when only one subunit is present.

Unlike Trf4 and Mtr4 RecA1, tRNA-binding sites in other Mtr4 domains and Air2 showed altered D-uptake between tRNA-bound Mtr4 or Trf4-Air2 and tRNA-bound TRAMP. Other Mtr4 domains involved in recognizing tRNA are the RecA2 and the Fist (26). In all cases, these domains had reduced D-uptake upon adding tRNA, but the reduction was slightly less in TRAMP than in Mtr4 alone (**Fig. 1B, E, F** and **Fig. 2B, D**). This difference suggests a slight decrease in the occupancy of these binding sites in TRAMP, as opposed to a gross change in the tRNA-binding path. We concomitantly observed the opposite trend for Air2 tRNA-binding motifs, ZnKs 2-3. These ZnKs had reduced D-uptake upon adding tRNA, but the reduction was slightly more in TRAMP than Trf4-Air2 (**Fig. 3B, D**). These data suggest that within TRAMP, these RNA-binding regions of Mtr4 and Air2 are competitive and that in our system, the tRNA slightly prefers binding Air2 ZnKs 2-3 over Mtr4 RecA2 and Fist. The reduced D-uptake observed at late time points for regions of Trf4 and Air2 ZnK 5 is also consistent with increased tRNA occupancy of Trf4-Air2 within TRAMP (**Fig. 3B, D**, and **Fig. S4F**).

A unifying explanation for the HDX results is that TRAMP has various tRNA-binding modes. These include (1) tRNA binding Mtr4 RecA1, RecA2, and Fist, just as in Mtr4, (2) tRNA binding to the positively charged surface formed by the Trf4 catalytic domain and NRM, and Air2 ZnKs 2-3, and (3) tRNA binding to Mtr4 and Trf4-Air2 simultaneously in a hybrid manner (**Fig. 5**). The hybrid state involves Mtr4 RecA1 and Air2 ZnKs 2-3 and may or may not involve the Trf4 catalytic domain and NRM. Given that the Mtr4 RecA2 and Air2 ZnKs 2-3 appear competitive for tRNA binding, we hypothesize that the hybrid state is an intermediate in the transfer of tRNA between the other two states. However, it could theoretically be an initial free tRNA binding state. If it is an intermediate, it suggests that tRNA transfer between active sites occurs on the relatively slow HDX time scale (minutes to hours). tRNA transfer may accelerate with a more wild-type poly(A) 3’ tail (we have three non-A bases on the 3’ end) or ATP turnover. We predict that the nature of the RNA substrate will also influence the prevalence of each binding mode.

**Figure 5.**
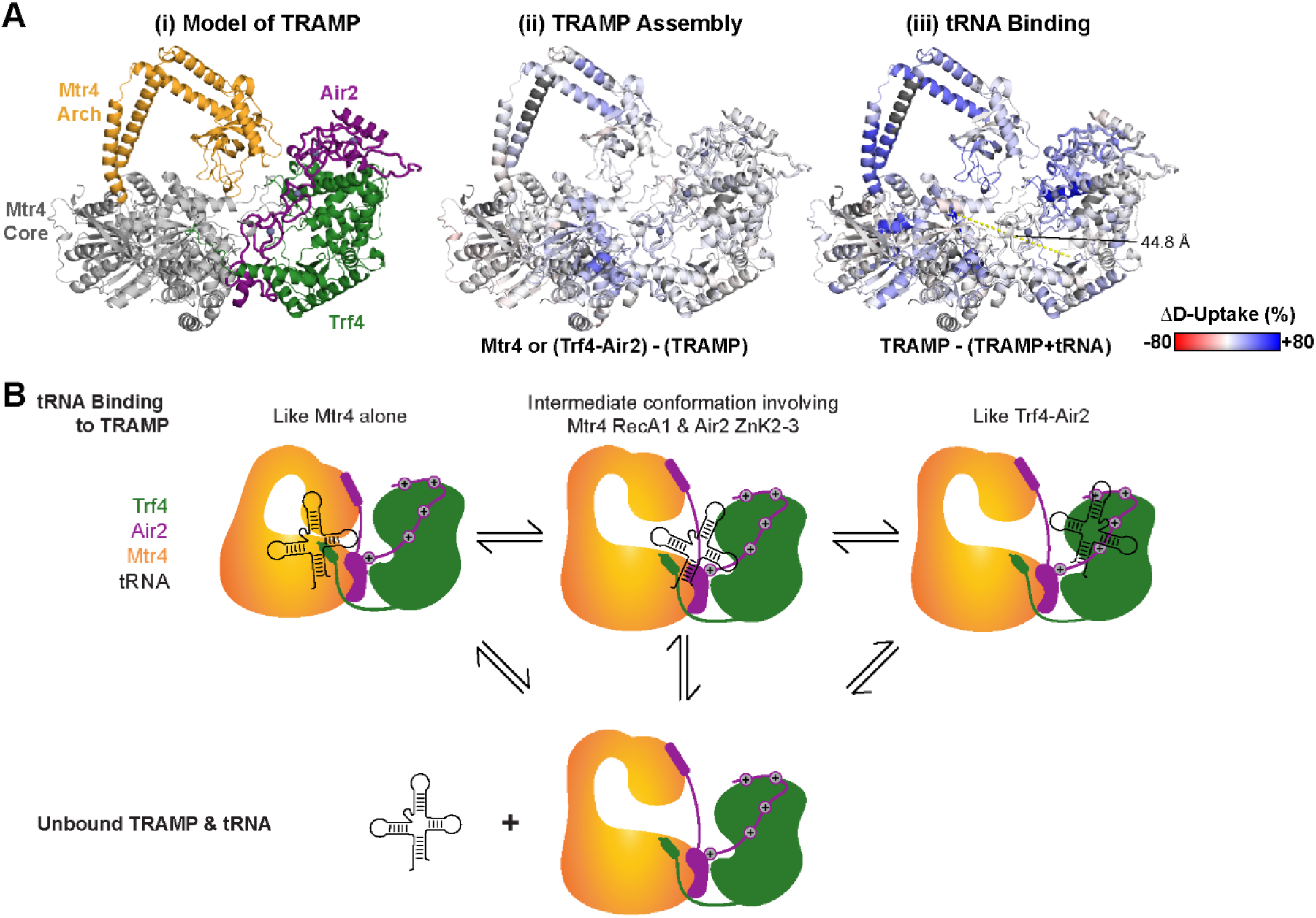
Model of TRAMP assembly and tRNA binding. (A) Regions of the Alphafold2 Multimer model of TRAMP with a pLDDT score > 40 (see **Fig. S5A**). The model also shows lower confidence (pLDDT ≤ 40), connecting regions of Trf4 between residues 128-155 for continuity with the Mtr4 Interaction site (residues 120-127). In (**i**), Mtr4 Arch is orange, Mtr4 helicase core is gray, Trf4 is green, Air2 is purple, and zinc atoms are spheres. In (**ii**), the model is colored by the difference in Mtr4 or Trf4-Air2 D-uptake upon TRAMP assembly. In (**iii**), the model is colored by the difference in TRAMP D-uptake upon tRNA binding. For (**ii**) and (**iii**), the differences are the percent change in D-uptake per residue summed across all time points. The scale is -80 to 80% (red to white to blue) based on DynamX residue-level scripts without statistical filtering. Residues without coverage are gray, and zinc atoms are spheres. (**B**) Cartoon depicting various tRNA-bound states of TRAMP, including an intermediate where tRNA is simultaneously bound to Mtr4 RecA1 (but not RecA2 or the Fist) and Air2 ZnK2 and ZnK3. This intermediate may represent a handoff between Trf4-Air2 and Mtr4.

Beyond tRNA-binding domains, we also observed different D-uptake signatures for the Mtr4 Arch and Air2 residues 37-45 in tRNA-bound Mtr4 or Trf4-Air2 and tRNA-bound TRAMP (**Fig. 2B, D**, and **Fig. 3B, D**). Within the Mtr4 Arch, helix 1 mimicked trends seen for the Fist, with slightly more D-uptake in tRNA-bound TRAMP than tRNA-bound Mtr4, consistent with the Arch being less constrained due to less tRNA occupancy at the Fist. Helix 2 and helix 3 trends are convoluted as TRAMP assembly and tRNA binding influence their D-uptake (**Fig. 2D**), but the fact that they have different D-uptake between tRNA-bound TRAMP and tRNA-bound Mtr4 suggests different Arch dynamics. Air2 residues 37-45, which interface with the Mtr4 helicase core, also had slightly less D-uptake in tRNA-bound TRAMP than in tRNA-bound Mtr4 (**Fig. 3D**). These differences reflect subtle conformation changes that occur only within TRAMP upon interaction with tRNA and reinforce crosstalk between Mtr4 and Trf4-Air2.

To further investigate the idea of RNA transfer between Mtr4 and Trf4-Air2, we used AlphaFold2 Multimer to generate TRAMP models with full-length proteins. The 25 models had average pLDDT scores between 0.57 and 0.68; the highest-scoring model is shown (**Fig. 5A** and **Fig. S5A**). Most models had Trf4-Air2 interactions with Mtr4, as seen in the crystal structure (31), although none of the models had the Air2 AIM near the Mtr4 Fist. To accommodate interaction between the Air2 AIM and Mtr4 Fist, the Mtr4 Arch must reposition upon TRAMP assembly, as shown by our HDX data. All models place the Trf4-Air2 and Mtr4 active sites side-by-side, also consistent with our HDX data as regions with decreased D-uptake upon TRAMP assembly map along a central plane (**Fig. 5A**). Further, all models have an unobstructed pathway between the Trf4-Air2 polymerase active site and Mtr4 helicase site, allowing for potential transfer of an RNA substrate (**Fig. S5B**). The two active sites are approximately 45 Å apart, suggesting that peripheral RNA-binding domains may mediate RNA transfer. Our observed competition between the Mtr4 Fist and Air2 ZnKs 2-3 suggests these may be key to this process, at least for tRNA. The conformational variation of the Mtr4 Arch and inherent flexibility of Air2, evident from high-resolution structural work and HDX, make them suited to the dynamic activity of RNA transfer. We provide a model for RNA transfer between Trf4-Air2 and Mtr4, revealing multiple tRNA-binding modes and demonstrating competition between distinct RNA-binding motifs within TRAMP. Our experiments and modeling highlight the dynamic nature of TRAMP, revealing a complex network of interfaces that may facilitate RNA transfer between Trf4-Air2 and Mtr4.

## Discussion

Our solution studies reaffirm previously seen interfaces between Mtr4 and Trf4-Air2 (31) and reveal substantial conformational rearrangement of the Mtr4 Arch and Air2 ZnKs upon TRAMP assembly. The significantly reduced exchange within the Mtr4 Arch shows that Mtr4 more frequently samples a closed conformation in TRAMP than in free Mtr4 (**Fig. 2E**). Mtr4 closed versus open conformations have been discussed previously (21, 26), with the closed conformation characterized by reduced dynamics of the Arch helical stalk and intra-chain contacts between the Fist and surface of the RecA2. The Mtr4 closed conformation induced by TRAMP assembly resembles that in Mtr4 bound by tRNA, although tRNA has a more pronounced effect. Direct tRNA contacts with the Fist are responsible for the closed conformation of tRNA-bound Mtr4 (26), and such contacts may be mimicked by the Air2 AIM within TRAMP.

While AIMs from other proteins, such as Nop53, Utp18, and NVL (27, 28, 44), play a critical role in binding to Mtr4, the Air2 AIM appears dispensable for TRAMP assembly and unwinding activity. The Air2 AIM only transiently interacts with the Mtr4 Fist and does not contribute significantly to the affinity between Trf4-Air2 and Mtr4. It seems likely that the Mtr4 closed conformation induced by TRAMP assembly through high-affinity interactions, such as Air2 binding on the Mtr4 helicase core, facilitates secondary, low-affinity interactions, such as those between the Air2 AIM and Mtr4 Fist. Distance constraints, in fact, preclude the Air2 AIM from binding the Fist if Air2 is simultaneously bound on the helicase core and Mtr4 is in an open conformation. A role for the Air2 AIM may emerge with different RNA substrates, polyadenylation states, or in polymerase activity assays. Transient binding of the Air2 AIM by the Mtr4 Fist may also prevent Mtr4 interactions with other AIM-containing proteins in vivo, similar to what was proposed for human exosome inhibitor NRDE2 (45). The Mtr4 Fist, on the other hand, contributes to the affinity between Trf4-Air2 and Mtr4 (**Fig. 3C**), suggesting a role beyond interaction with the Air2 AIM. No evidence exists for non-AIM interactions between Trf4-Air2 and the Mtr4 Fist or helical stalk. We suspect that removal of the Fist alters Arch conformational dynamics, thereby indirectly hindering TRAMP assembly.

TRAMP assembly and tRNA interaction also induce constraint in Air2 ZnKs in the context of the Trf4-Air2 sub-complex. The precise position and conformation of Air2 ZnKs 1-3 are challenging to pinpoint as they are dynamic in solution. Air2 ZnKs 1-3 fold cooperatively and become constrained by TRAMP assembly (**Fig. 3A**). Given that the region N-terminal to the Air2 ZnKs tightly binds the Mtr4 helicase core, this constraint likely modulates the distance between Trf4 and Mtr4 active sites within TRAMP, potentially playing a role in RNA transfer. Prior work has shown that Air2 ZnKs 2-3 bind tRNA in Air2 alone (32), and we demonstrate the maintenance of this function in the Trf4-Air2 complex (**Fig. 3B**). Modeling of Trf4-Air2 places Air2 ZnKs 2-3 alongside the Trf4 catalytic domain and NRM to create a continuous, positively charged channel primed for RNA recognition (**Fig. 4D**). The significantly reduced exchange of this channel upon adding tRNA affirms its role in RNA binding (**Fig. 4C**). The cleft between the catalytic domain and NRM of Trf4 is likely to recognize the 3’ end of the tRNA, similar to homologous polymerases (33, 34). There is an intriguing parallel between the Mtr4 Arch and the Air2 ZnKs as both undergo conformational constraint upon TRAMP assembly and contain motifs that directly bind RNA, although not the RNA 3’ end.

A comparison between tRNA binding to Mtr4 or Trf4-Air2 and TRAMP highlights essential aspects of TRAMP tRNA recognition. There was virtually no difference in tRNA binding to the Mtr4 RecA1 (**Fig. 1B**) or the cleft between the Trf4 catalytic and central domains (**Fig. S4F**), which are the motifs thought to recognize the RNA poly(A) tail or 3’ end. These motifs may bind tRNA simultaneously, or tRNA may rapidly transfer between them rather than contribute to the free RNA pool. At the same time, we observed competition between other tRNA binding motifs with increased exchange in the Mtr4 RecA2 and Fist (**Fig. 1B, and Fig. 2B**) offset by decreased exchange in Air2 ZnKs 2-3 (**Fig. 3B**). Since these changes were moderate, they likely represent a low-abundance population where the tRNA is bound to TRAMP in a hybrid manner involving the Mtr4 RecA1 and Air2 ZnKs 2-3, but not the Mtr4 RecA2 and Fist (**Fig. 5B**). This hybrid complex may be an intermediate in the transfer of tRNA between the more canonical tRNA-bound Mtr4 and Trf4-Air2 sub-complexes within TRAMP. We predict that ATP turnover and RNA features, such as 3’ poly(A) tail length, will influence the distribution between various tRNA binding modes of TRAMP.

Our experiments solely focus on one combination of TRAMP homologs found in *S. cerevisiae* (Trf4, Air2, and Mtr4). We anticipate that our results similarly apply to the other TRAMP homologs. Trf4 and Trf5 share 68% sequence identity and 80% similarity, while Air1 and Air2 exhibit 50% identity and 64% similarity overall. All Mtr4-interacting regions in Trf4 and Air2 are conserved in Trf5 and Air1(**Fig. S5C**). Specifically, Trf4_120-127_ and Trf5_104-111_ are highly conserved and contain the key aspartate and phenylalanine residues required for TRAMP complex formation (31). The N-terminal region of Air1/2 is also highly conserved, including the AIM sequence and the Mtr4 core interacting region. A four-residue insertion occurs between the AIM and Mtr4 core interacting region in Air1. This increased length could alter Arch dynamics between the two homologs and provide a mechanism for dealing with different RNA substrate preferences.

Our study underscores the dynamic nature of the TRAMP complex in *S. cerevisiae*, which we expect to be typical of TRAMP complexes across species. The interfaces between Mtr4 and Trf4- Air2 are small and sometimes involve disorder, and TRAMP assembly involves conformational changes in multiple domains. While the data presented here show the general position and orientation of the core domains of TRAMP, we anticipate that the precise positioning of Mtr4, Trf4, and Air2 is highly dynamic, especially in the absence of an RNA. We also expect TRAMP assembly and dynamics to be sensitive to specific characteristics of the RNA substrate. Similar dynamics may be at play in other Mtr4 complexes. A common theme appears to be the stabilization of short, low-complexity regions upon complex formation, as described in TRAMP, MTREC (46), NEXT (30), and exosome (22, 47) complexes. These interactions likely foster increased mobility in the relative positioning of individual components, possibly enabling engagement with a variety of substrates and co-factors. Complete description of these inherently dynamic systems will require continued studies using a combination of solution and structural approaches.

## Methods

### Protein Purification

*S. cerevisiae* (Sc) Mtr4 (P47047) was recombinantly expressed with an N-terminal, TEV-protease cleavable, hexahistidine tag in an *Escherichia coli* BL21-CodonPlus (DE3) cell line and purified as previously described (48). In brief, following cell lysis by sonication, Mtr4 was purified by Ni-NTA immobilized metal affinity, heparin affinity, ion exchange, and size exclusion chromatographies. Full-length ScTrf4 (P53632) and its point mutation variant (D236A/D238A), in which two aspartate residues in the active site were mutated to alanines, were used. These were co-expressed with ScAir2 residues 1-223 (Q12476) in an *E. coli* BL21-CodonPlus (DE3) cell line. Air2 was tagged with an N-terminal hexahistidine tag. Zinc was added to the culture during expression. The protein complex was purified as previously described (48). In brief, following cell lysis by sonication, Trf4-Air2 was purified by Ni-NTA immobilized metal affinity, heparin affinity, reverse-phase, and size exclusion chromatography. Additional Air2 and Mtr4 mutants (Mtr4^ΔFist^ Trf4- Air2^NΔ17^, and Trf4-Air2^ZKO^) were prepared using the same expression and purification protocol. Air2^NΔ17^ had the first 17 residues deleted, and Air2^ZKO^ had two point mutations, C84S and C104S. The Mtr4^ΔFist^ was constructed as previously described (26) by removing residues 667-813 and inserting a three-glycine linker using the Q5 Site-Directed Mutagenesis Kit (NEB).

### RNA Preparation

The tRNA _i_ ^Met^, dsRNA^16^, and dsRNA^32^ substrates used in this study are the same as previously described (26). The tRNA _i_ ^Met^ (transfer RNA initiator methionine) substrate is like the native tRNA _i_ ^Met^ sequence from *S. cerevisiae* with a CCA cap and non-native 5’ and 3’ cloning artifacts. It was generated using in vitro transcription with T7 RNA polymerase (ThermoFisher Scientific) on a plasmid linearized after the tRNA _i_ ^Met^ sequence, and purified under non-denaturing conditions as previously described (49). Briefly, the transcription reaction was loaded onto HiTrap DEAE Sepharose FF (GE), and the tRNA _i_ ^Met^ eluted off using a NaCl gradient. The dsRNA^16^ (double-stranded RNA, 16 base pairs) and dsRNA^32^ (double-stranded RNA hairpin with 32 base pairs) both contain a 5-adenosine poly(A) tail. These dsRNA substrates were purchased from Integrated DNA Technologies. The purchased oligonucleotides were annealed following the manufacturer’s protocol. Single strand RNA oligos were resuspended in Annealing Buffer (100 mM potassium acetate, 30 mM HEPES, pH 7.5), mixed in equal molar amounts, heated to 94°C for 2 min, and then allowed to cool to room temperature by turning off the heating block. Annealed RNAs were then diluted to 10 µM, flash frozen with LN_2_, and stored at -80ºC until use.

### Hydrogen-Deuterium Exchange with Mass Spectrometry

HDX reactions were prepared from stocks of 12.5 µM Mtr4, Trf4_D236A+D238A_-Air2_1-223_, and Mtr4- Trf4_D236A+D238A_-Air2_1-223_ each with or without 25 µM tRNA _i_ ^Met^ (i.e. 2-fold molar excess). For analysis of Air2^ZKO^, HDX reactions were prepared from stocks of 12.5 µM Mtr4, Trf4_D236A+D238A_-Air2_1-223_, Trf4_D236A+D238A_-Air2_1-223_^ZKO^, and Mtr4-Trf4^D236A/D238A^-Air2_1-223_^ZKO^. In all cases, stocks were in 50 mM HEPES, 100 mM NaCl, 0.5 mM MgCl_2_, 0.2 mM TCEP, pH 6.5, and incubated for at least 10 min at 25°C. Samples were diluted 2:23 with the same buffer containing H_2_O for controls or D_2_O for exchange samples (pH_read_=6.1 at 25°C and final D_2_O of 92%). Exchange proceeded at 25°C for 10, 10^2^, 10^3^, 10^4^, or 10^5^ s for the wild-type experiment, and 10, 10^2^, or 10^3^ s for the Air2^ZKO^ experiment. Exchange was quenched by mixing samples 1:1 (v:v) with cooled 1% (v/v) formic acid, 3.84 M guanidinium chloride-HCl, pH 1.75 (mix had a final pH of 2.3 at 0°C), and samples were flash frozen in LN_2_. Samples were prepared in triplicate and stored at −80°C. Samples were thawed for 50 s immediately prior to injection into a Waters™ HDX manager in line with a SYNAPT G2-Si. In the HDX manager, samples were digested by *Sus scrofa* Pepsin A (Waters™ Enzymate BEH) at 15°C and the peptides trapped on a C4 pre-column (Waters™ Acquity UPLC Protein BEH C4) at 1°C using a flowrate of 100 µL/min for 3 min. The chromatography buffer was 0.1% (v/v) formic acid. Peptides were then separated over a C18 column (Waters™ Acquity UPLC BEH) at 1°C and eluted with a linear 3–40% (v/v) acetonitrile gradient using a flowrate of 40 µL/min for 7 min. Samples were injected in a random order.

Mass spectrometry data were acquired using positive ion mode in either HDMS or HDMS^E^ mode. Peptide identification of water-only control samples was performed using data-independent acquisition in HDMS^E^ mode. Peptide precursor and fragment data were collected via collision-induced dissociation at low (6 V) and high (ramping 22–44 V) energy. HDMS mode was used to collect low-energy ion data for all deuterated samples. All samples were acquired in resolution mode. Capillary voltage was set to 2.8 kV for the sample sprayer. Desolvation gas was set to 650 L/h at 175°C. The source temperature was set to 80°C. Cone and nebulizer gas was flowed at 90 L/h and 6.5 bar, respectively. The sampling cone and source offset were both set to 30 V. Data were acquired at a scan time of 0.4 s with a range of 100–2000 m/z. Mass correction was done using [Glu1]-fibrinopeptide B as a reference. For ion mobility, the wave velocity was 700 ms^−1,^ and the wave height was 40 V. Peptide lists were generated from water-only controls in PLGS (Waters™ Protein Lynx Global Server 3.0.2) using a database containing S. scrofa Pepsin A and *S. cerevisiae* Mtr4, Trf4_D236A+D238A_ and Air2_1-223_ or Air2_1-223_^ZKO^. In PLGS, the minimum fragment ion matches per peptide was 3, and methionine oxidation was allowed. The low and elevated energy thresholds were 250 and 50 counts, respectively, and the overall intensity threshold was 750 counts. DynamX 3.0 was used to search the deuterated samples for peptides with 0.3 products per amino acid, and one consecutive product found in 2 out of 7–10 controls. Peptide assignments were manually validated.

Significance thresholds were calculated for each protein following the method of Hageman and Weis (50). As the computed significance thresholds for Mtr4 and Trf4 ranged between 0.22 and 0.39 Da, a 0.4 Da cutoff was used for these proteins in both data sets. Calculated significance thresholds for Air2 ranged between 0.45 and 0.50 Da, so a 0.5 Da cutoff was used for Air2 across both data sets. Back-exchange was calculated using peptides from the TRAMP sample that had plateaued and had a D-uptake >40%. Structural images were made using scripts from DynamX in the PyMOL Molecular Graphics System, Version 2.5 (Schrödinger, LLC), and heat maps were made using HD-eXplosion (51). The mass spectrometry proteomics data have been deposited to the ProteomeXchange Consortium via the PRIDE partner repository with the dataset identifier PXD053773 (52). The summary table and centroids are included in **Table S1**.

### RNA Unwinding Assays

Pre-steady state unwinding activity was determined by monitoring the displacement of a 16-nt RNA labeled with fluorescein on its 5’ end from a longer unlabeled RNA strand when incubated with TRAMP at saturating levels of dATP. Unwinding reactions were performed in 40 mM MOPS, 100 mM NaCl, 0.5 mM MgCl_2_, 5% (v/v) glycerol, 0.01% (v/v) nonidet-P40 substitute, 1 U/μL of Ribolock (ThermoFisher Scientific), 2 mM DTT, pH 6.5 at 30°C. 400 nM TRAMP was reconstituted at 500 nM from independently purified Mtr4 and Trf4-Air2 and incubated for 10 min at 30°C. After incubation, the reconstituted TRAMP with 10 nM fluorescein-labeled duplex RNA and 100 nM DNA-trap oligonucleotide for 10 min at 30°C. Reactions were initiated by the addition of 1.6 mM dATP with MgCl_2_. At various times, aliquots were quenched with a 1:1 dilution in 100 mM EDTA, 50 mM Tris-HCl, 4 mg/ml Proteinase K, 20% (v/v) glycerol at pH 8.0. Samples were separated on a 15% native polyacrylamide TBE gel for 80 min at 120 V. Gels were imaged with a ChemiDoc MP Imaging system (Bio-Rad).

### Surface Plasmon Resonance

Binding kinetics were performed using the OpenSPR instrument (Nicoya Lifesciences) using a Ni-NTA sensor chip maintained at 20ºC. All proteins were diluted substantially with SPR running buffer (10 mM HEPES, 150 mM NaCl, 3 mM EDTA, 5% (v/v) glycerol, 0.01% (v/v) Tween20, 100 μg/mL His-IpaD, pH 7.5) before the experiment. His-tagged Trf4-Air2, Trf4-Air2^NΔ17^, or Trf4- Air2^ZKO^ (diluted in running buffer without His-IpaD) at a concentration of 50 μg/ml was immobilized on the chip. Mtr4 or Mtr4^ΔFist^ was then injected as analytes at concentrations varying from 62.5 nM to 5 μM. The flow rate was maintained at 20 μL per minute, and the buffer was flowed over the chip for an additional 5 min after terminating the flow of analyte. After about 10 min and the SPR response returned to baseline, the experiment was repeated at different analyte concentrations. SPR data was analyzed using TraceDrawer software (Nicoya Lifesciences). All K_D_ values were calculated using 4 to 5 concentrations per analyte and each experiment repeated in triplicate, with a new SPR chip each time, with at least one biological replicate in the set.

### Size Exclusion Chromatography

TRAMP complex formation analysis was performed using a 24 mL Superdex 200 Increase 10/300 GL. Mtr4 and Trf4-Air2 were mixed in a 1:1 ratio and diluted to 10 µM in running buffer (50 mM HEPES pH 7.5, 160 mM NaCl, 5%(v/v) glycerol, 2 mM BME) and incubated on ice for 30 min. The different TRAMP mutants were then injected over a column previously equilibrated with running buffer. Eluted protein fractions were monitored using absorbance at 280 nm and analyzed by 12% acrylamide SDS-PAGE.

## Acknowledgments

This work was supported by the National Institute of General Medical Sciences, NIH (R01GM117311 and R15GM148949 to S.J. and R35GM133751 to S.D). We thank Mark Gold and Kami Morgan for their assistance with mutagenesis and members of the D’Arcy and Johnson Groups for scientific discussions.

