## Supplement for "TRAMP assembly alters the conformation and RNA binding of Mtr4 and Trf4-Air2"

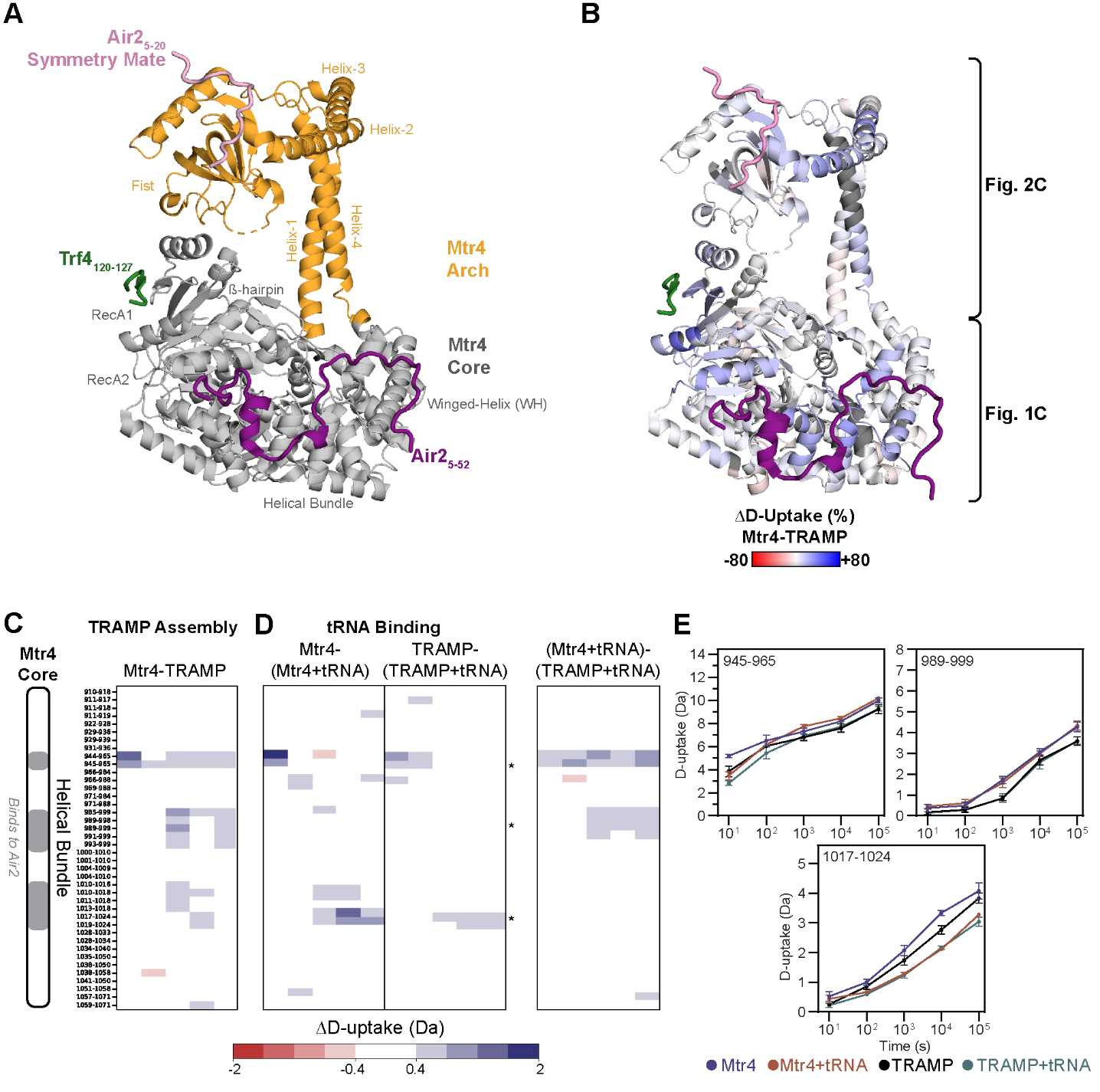

**Figure S1. Changes in D-uptake of Mtr4 Helical Bundle upon TRAMP assembly and tRNA binding**

(**A**) TRAMP structure where Mtr4 Arch is orange, Mtr4 Core is gray, Trf4 is green, Air2 is purple, and Air2 from a symmetry-related Mtr4 is pink. The structure is PDB 4U4C. (**B**) TRAMP structure as in **A**, except Mtr4 is colored by the percent difference in D-uptake per residue between Mtr4 and TRAMP summed across all time points. The scale is -80 to 80% (red to white to blue) based on DynamX residue-level scripts without statistical filtering. Mtr4 residues without coverage are gray. (**C**) The difference in Mtr4 Helical Bundle D-uptake upon TRAMP assembly. (**D**) The difference in Mtr4 Helical Bundle D-uptake upon tRNA binding to Mtr4 (left) or TRAMP (middle); direct comparison of the two tRNA-bound complexes (right). For **C** and **D**, red/blue indicates a difference of ≥|0.4| Da and a *p*-value ≤0.01 in a one-sided Welch’s t-test (n=3). Exchange time points were 10^1^-10^5^ s (y-axis). (**E**) Select D-uptake plots of Mtr4 Helical Bundle peptides showing Mtr4 (blue), Mtr4 with tRNA (orange), TRAMP (black), and TRAMP with tRNA (green). Error bars are ±2SD of the average with n=3. The y-axis is 80% of the maximum theoretical D-uptake, assuming complete back exchange of the N-terminal residue of each peptide.

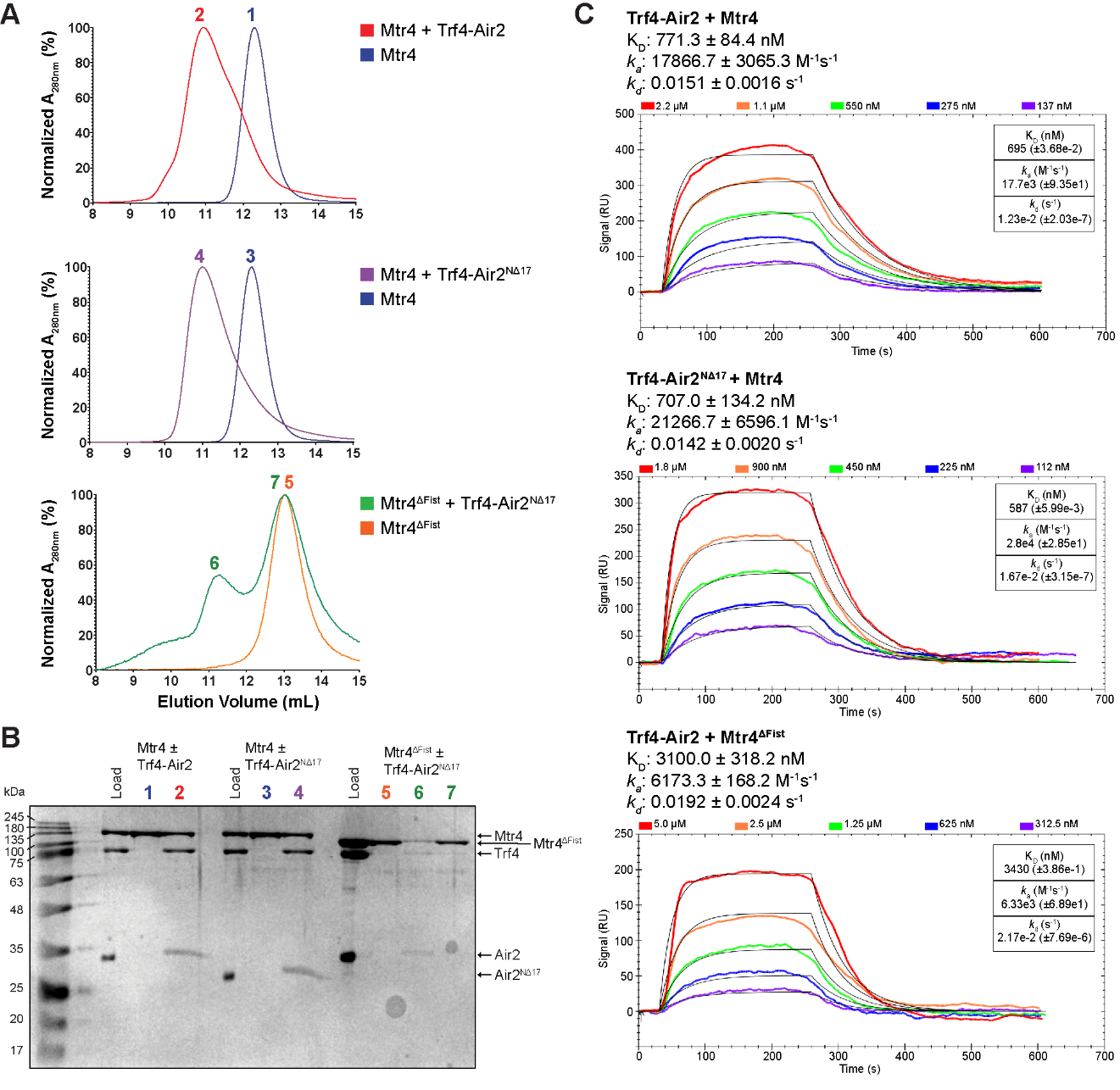

**Figure S2. SEC and SPR Analysis of TRAMP assembly**

(**A**) SEC of 10 μM Mtr4 and TRAMP (equimolar Mtr4 and Trf4-Air2) with wild-type proteins, Trf4-Air2^NΔ17^, and/or Mtr4^ΔFist^. The column was a Superdex 200 Increase10/300 GL equilibrated in 50 mM HEPES pH 7.5, 160 mM NaCl, 5% (v/v) glycerol, and 2 mM BME. (**B**) 12% SDS-PAGE of peak fractions from **A**. Peak and lane labels match. Gel was Coomassie stained. (**C**) Representative SPR response curves for TRAMP assembly with wild-type proteins, Trf4-Air2^NΔ17^, and/or Mtr4^ΔFist^. Trf4-Air2^WT or NΔ17^ was immobilized on a Ni-NTA sensor chip via an N-terminal hexahistidine tag on Air2. Mtr4^WT or ΔFist^ was injected at 20 μL/min at concentrations ranging from 62.5 nM to 5 μM. Graph insets show parameters from the global fitting of the single titration shown. Average parameters from at least 3 replicate titrations, including at least one biological replicate, are listed above the graphs and compared in **Fig. 3C**.

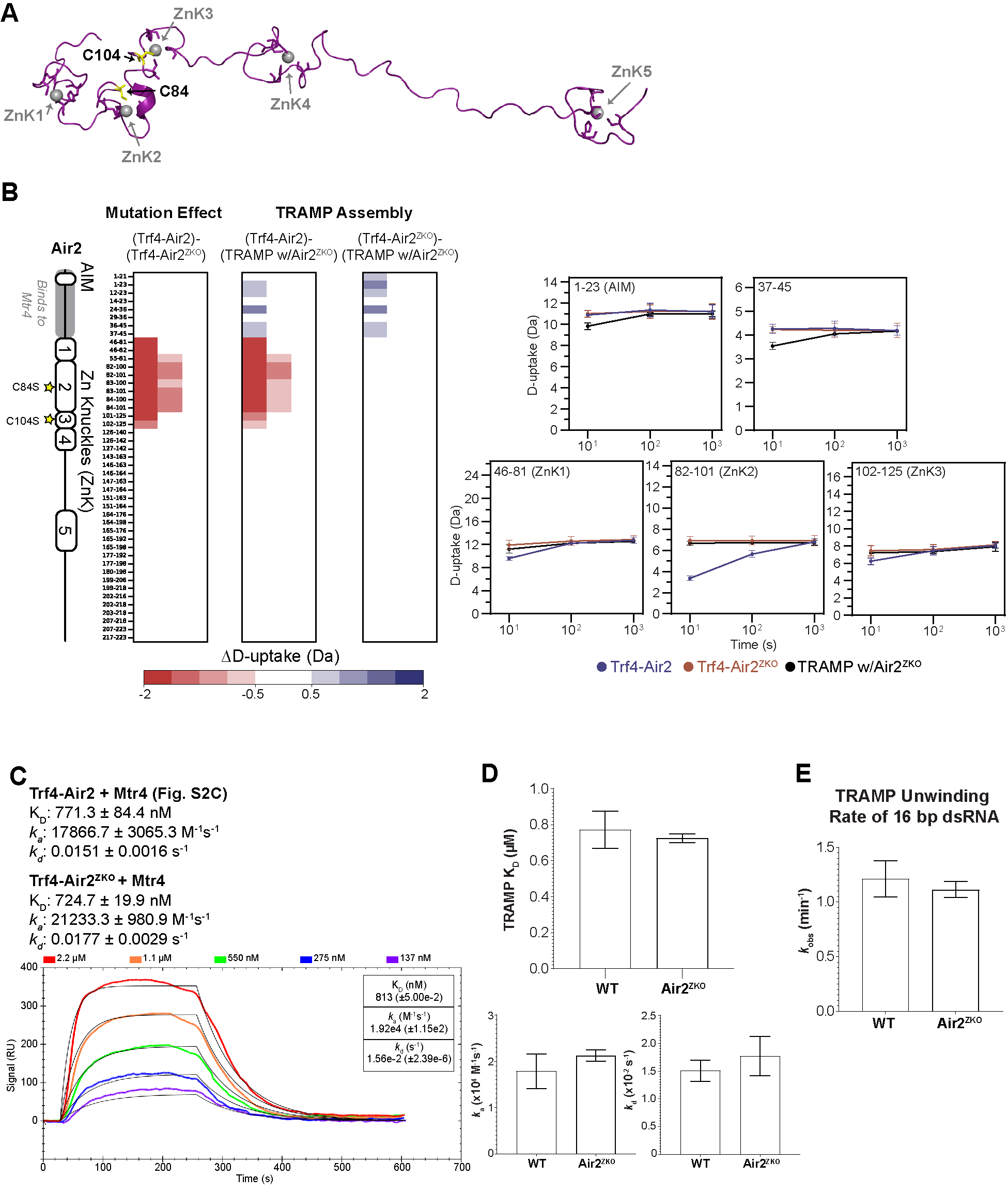

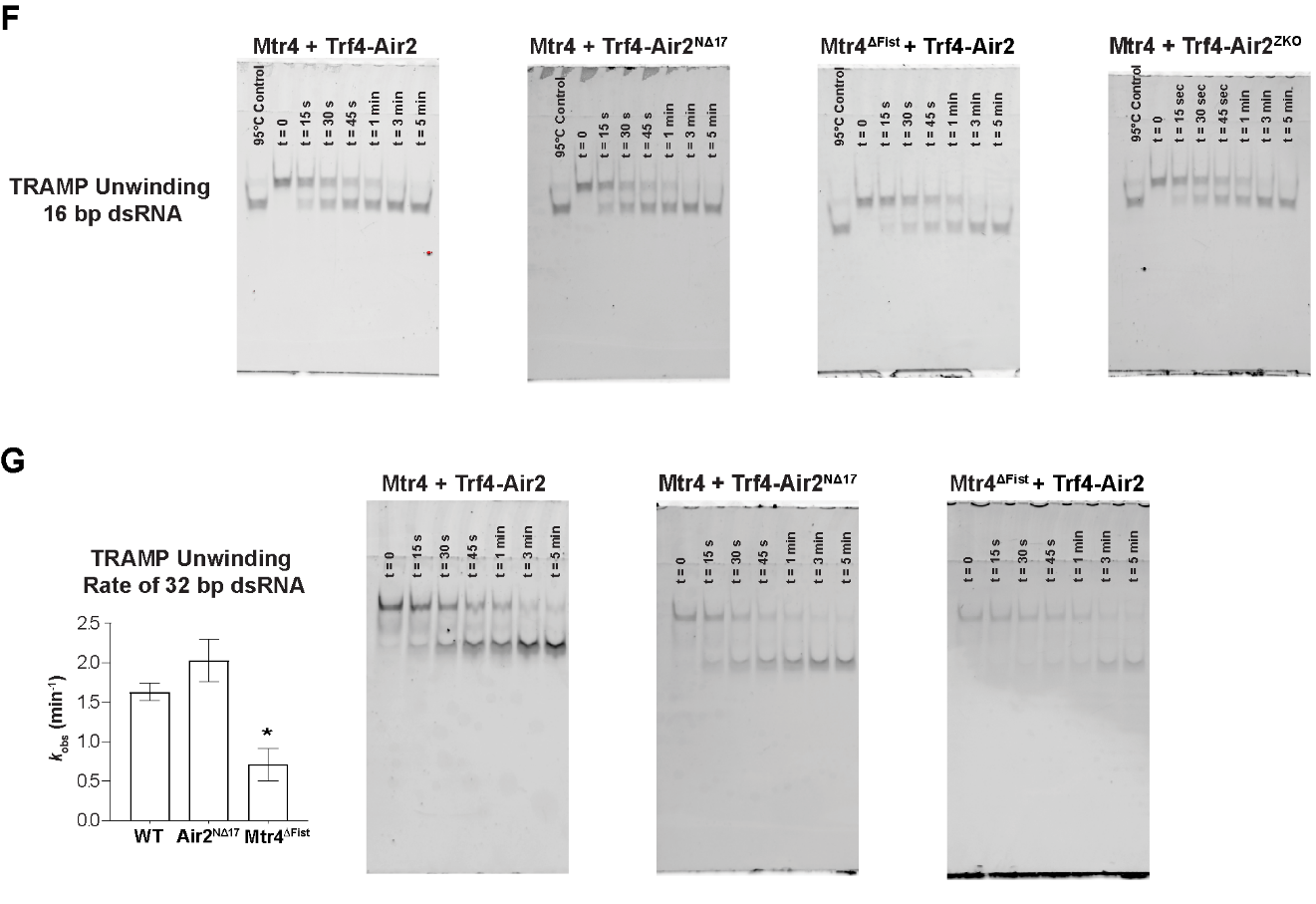

**Figure S3. Characterization of Trf4-Air2^ZKO^**

(**A**) Air2 structure where Air2 is purple, zinc atoms are spheres, and the mutated Cys84 and Cys104 residues are yellow sticks. The structure is PDB 2LLI. (**B**) The difference in Air2 D-uptake between Trf4-Air2 and Trf4-Air2^ZKO^ (left), Trf4-Air2 and TRAMP with Air2^ZKO^ (middle), and Trf4-Air2^ZKO^ and TRAMP with Air2^ZKO^ (right). Red/Blue indicates a difference ≥|0.5| Da and a *p*-value ≤0.01 in a one-sided Welch’s t-test (n=3). Exchange time points were 10^1^-10^5^ s (y-axis). The far-right shows select D-uptake plots of Air2 peptides for Trf4-Air2 (blue), Trf4-Air2^ZKO^ (orange), and TRAMP with Trf4-Air2^ZKO^ (black). Error bars are ±2SD of the average with n=3. The y-axis is 80% of the maximum theoretical D-uptake, assuming complete back exchange of the N-terminal residue of each peptide. (**C**) Representative SPR response curves for TRAMP assembly with Mtr4 and Trf4-Air2^ZKO^. Trf4-Air2^ZKO^ was immobilized on a Ni-NTA sensor chip via an N-terminal hexahistidine tag on Air2. Mtr4 was injected at 20 μL/min at concentrations ranging from 137 nM to 2.2 μM. Graph inset shows parameters from the global fitting of the single titration shown. Average parameters from at least 3 replicate titrations, including at least one biological replicate, are listed above the graph. (**D**) Comparison of TRAMP assembly parameters (K_D_, *k_a_*, and *k_d_*) measured by SPR. (**E**) Helicase activity using a synthetic 16 bp dsRNA substrate with a 5x adenosine 3’ overhang and 500 nM TRAMP. For **D** and **E**, TRAMP was wild-type (WT) (same data as in **Fig. 3C, E**), or with Air2^ZKO^. An asterisk (*) indicates a *p*-value ≤0.05 in a one-way ANOVA with the posthoc Tukey HSD test. Error bars show ±SD of the average with n=3, including at least 1 biological replicate. (**F**) Representative 15% native PAGE for measuring the unwinding of a 16 bp dsRNA by various TRAMP complexes. Bands were visualized and quantified using fluorescein on the 5’ end on the longer RNA strand. Quantified data are shown in **E** and **Fig. 3E.** (**G**) As for **E** and **F**, but with a synthetic 32 bp dsRNA substrate.

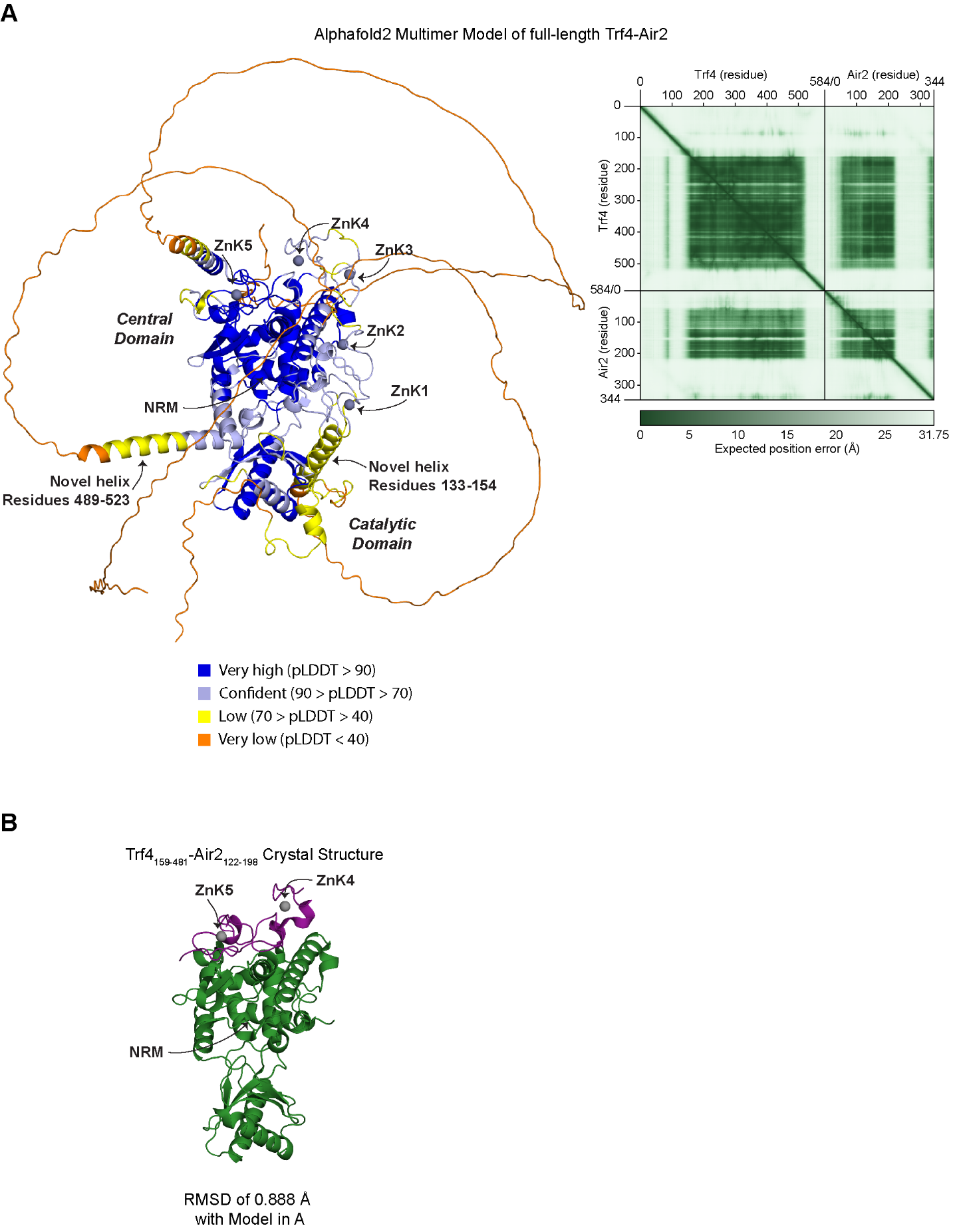

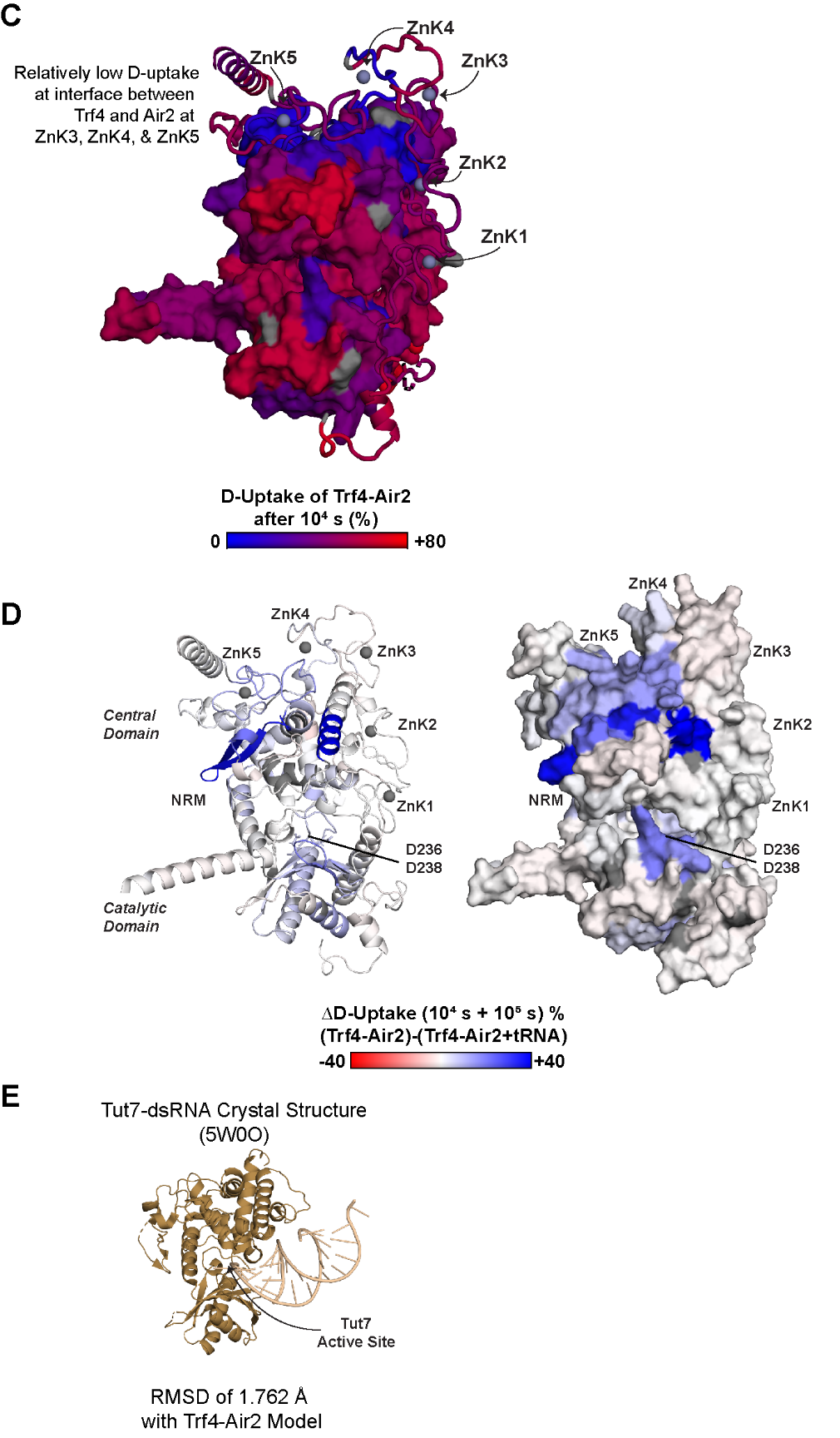

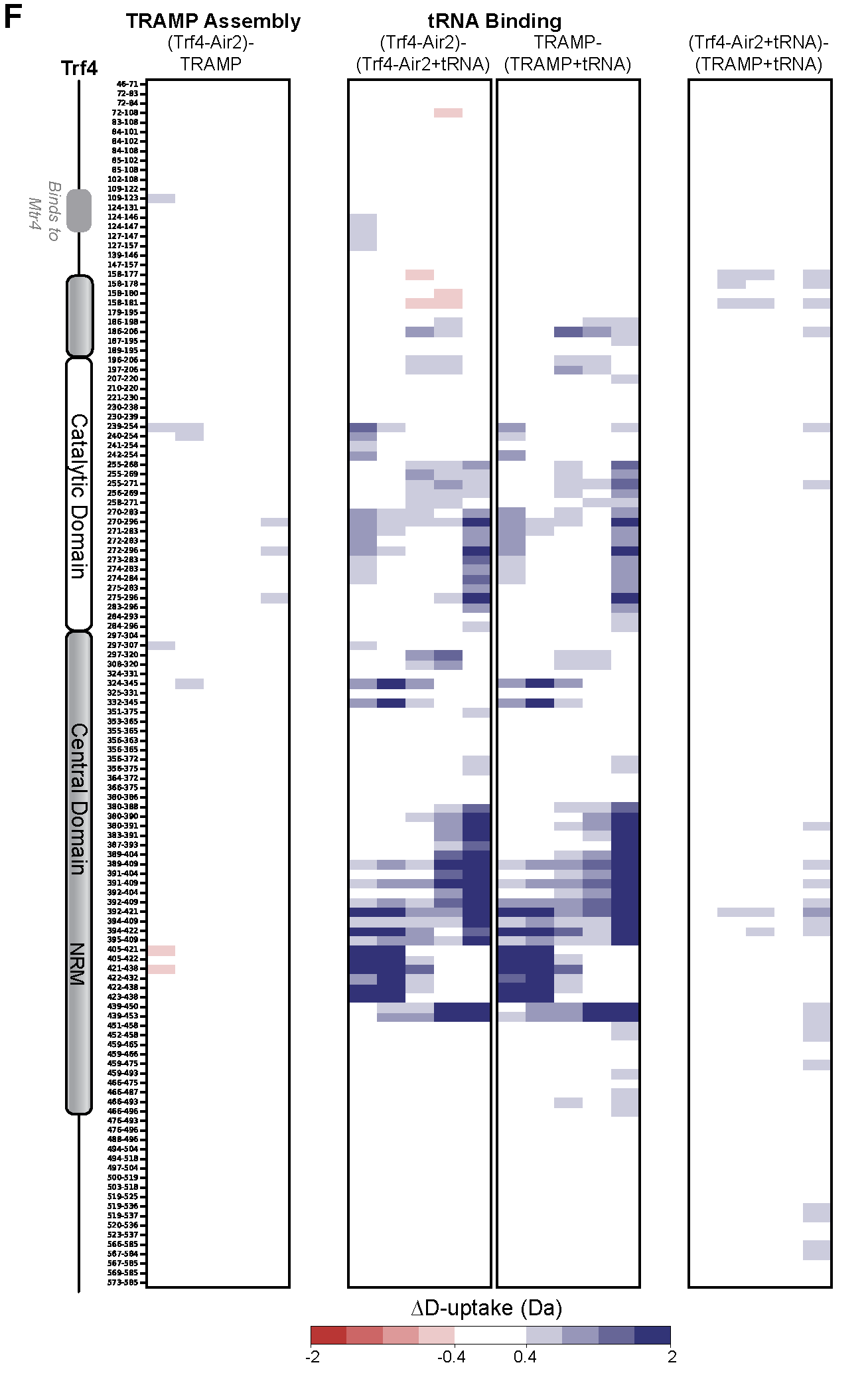

**Figure S4. Understanding Trf4 interaction with Air2 and tRNA**

(**A**) Alphafold2 Multimer Model of full-length Trf4-Air2. The model is colored by a position confidence score. Zinc atoms were placed by pairwise fitting ZnKs from two other Air2 structures (ZnK4 and ZnK5 from PDB 3NYB and ZnK1, ZnK2, and ZnK3 from PDB 2LLI). The top right shows the PAE (Predicted Aligned Error) confidence matrix for the model displaying paired per residue confidence. (**B**) Crystal structure of Trf4_159-481_-Air2_122-198_ (PDB 3NYB) aligned with the Alphafold2 Multimer model (RMSD of 0.888 Å). Trf4 is green, Air2 is purple, and zinc atoms are spheres. (**C**) Alphafold2 Multimer model of Trf4-Air2 colored by the percent D-uptake per residue after 10^4^ s exchange (Trf4 as surface and Air2 as cartoon). The scale is 0 to 80% (blue to red) based on DynamX residue-level scripts. Trf4-Air2 residues without coverage are gray, and zinc atoms are spheres. (**D**) Alphafold2 Multimer model of Trf4-Air2 colored by the percent change in D-uptake per residue upon adding tRNA summed across 10^4^ and 10^5^ s time points (left as cartoon, right as surface). The scale is -80 to 80% (red to white to blue) based on DynamX residue-level scripts without statistical filtering. Trf4-Air2 residues without coverage are gray, and zinc atoms are spheres. Zincs were placed in the model by pairwise superposition of coordinating residues in prior structures. Images contain only high-confidence regions (pLDDT > 40) which are Trf4_133-523_ and Air2_24-220_. (**E**) Structure of Trf4 homolog Tut7 bound to dsRNA (PDB 5W0O, RMSD of 1.762 Å). Tut7 is gold, and dsRNA is beige. (**F**) Expansion of **Fig. 4A-B**. The difference in Trf4 D-uptake upon TRAMP assembly (left); the difference in Trf4 D-uptake upon tRNA binding to Trf4-Air2 or TRAMP (middle two panels); direct comparison of the two tRNA-bound complexes (right). Red/Blue indicates a difference of ≥|0.4| Da and a *p*-value ≤0.01 in a one-sided Welch’s t-test (n=3). Exchange time points were 10^1^-10^5^ s (y-axis).

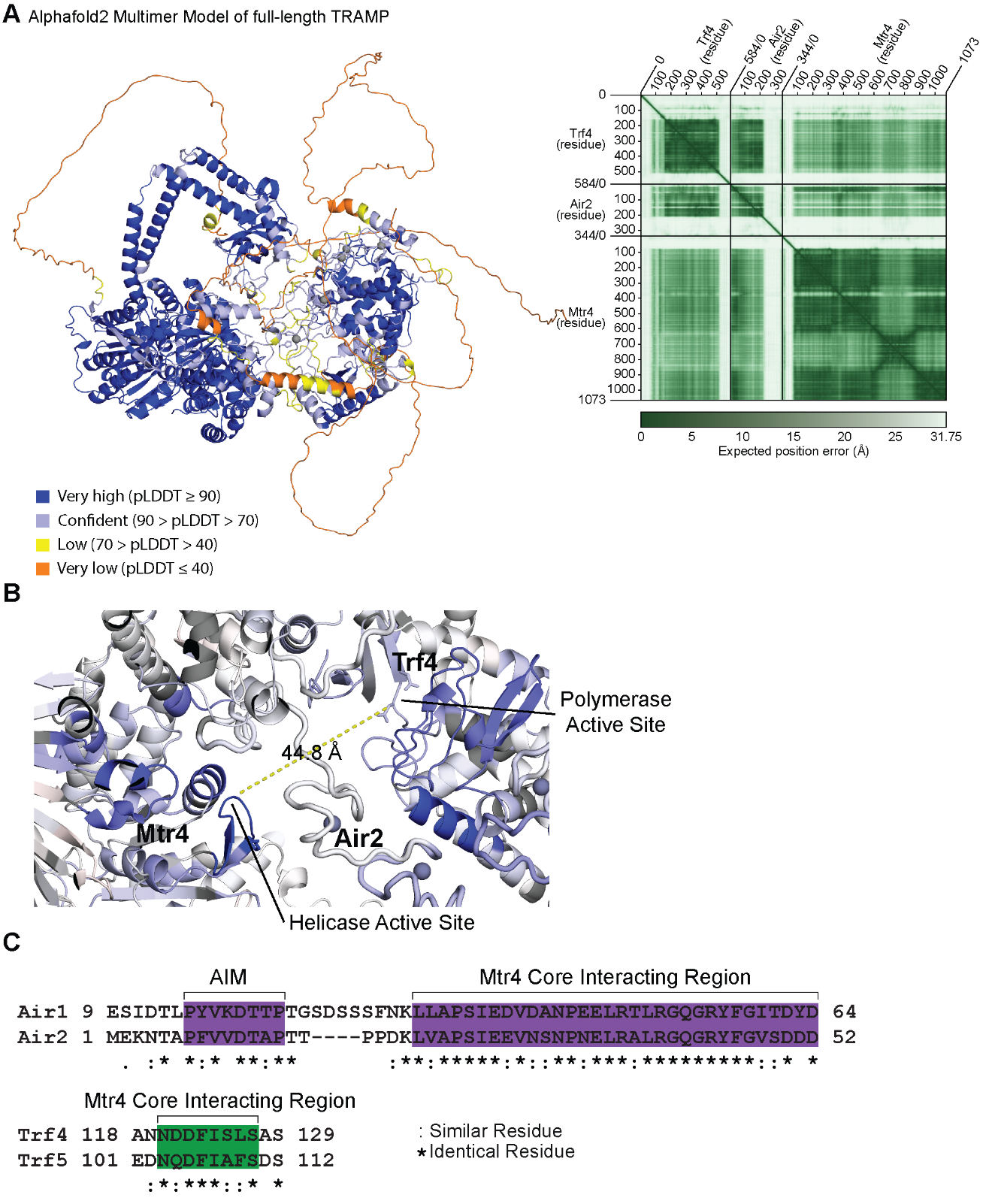

**Figure S5. Model of TRAMP**

(**A**) Alphafold2 Multimer Model of full-length TRAMP (Trf4-Air2 with Mtr4). The model is colored by a position confidence score. Zinc atoms were placed by pairwise fitting ZnKs from two other Air2 structures (ZnK4 and ZnK5 from PDB 3NYB and ZnK1, ZnK2, and ZnK3 from PDB 2LLI). The top right shows the PAE (Predicted Aligned Error) confidence matrix for the model displaying paired per residue confidence. (**B**) Zoom in on **Fig. 5A (iii)** to show the distance between active sites in TRAMP. (**C**) Sequence alignment of Mtr4 Core interacting regions of Air1 and Air2, and Trf4 and Trf5. The residues important at Air2-Mtr4 (purple) and Trf4-Mtr4 (green) interfaces are conserved between the homologs.
